# The Hidden Potential of PDE4 Inhibitor Rolipram: A Multifaceted Examination of its Inhibition of MMP2/9 Reveals Therapeutic Implications

**DOI:** 10.1101/2024.05.16.594542

**Authors:** Arka Bagchi, Analabha Roy, Anindya Halder, Arunima Biswas

**Author notes:** Corresponding authors: Analabha Roy: Assistant Professor, Department of Physics, The University of Burdwan, Bardhaman, West Bengal, India. PIN: 713104.,., Arunima Biswas: Assistant Professor, Cell and Molecular Biology Laboratory, Department of Zoology, University of Kalyani, Kalyani, Nadia, West Bengal, India. PIN: 741235.,.

## Abstract

A PDE4 inhibitor, Rolipram, was previously found to down-regulate (in a manner dependent on cAMP-PKA) MMP2 and MMP9 levels, important markers of epithelial-to-mesenchymal transition in human breast cancer cell lines. However, zymographic studies revealed that rolipram could also alter the enzymatic activities of these MMPs, even in the presence of the PKA inhibitor H89. This calls for more detailed investigations of the inhibitory mechanism of rolipram on MMP2 and MMP9. The prediction of ligand-based targets through online reverse screening indicated that proteases are likely targets of rolipram. Computational molecular docking also demonstrated significant binding affinities of rolipram for both MMP2 and MMP9 proteins. Concurrently, a well-known inhibitor of MMPs, SB3CT, was utilized as a positive control for this study. The best models of the docked complexes were used as initial conditions for molecular dynamics (MD) simulations to explore their dynamic behavior and stability. In particular, both the MMP2-rolipram and MMP9-rolipram complexes were found to be stable and compact for the duration of the simulation (300 ns). Several stable hydrogen bonds were also detected between the proteins and rolipram. *In vitro* experiments using primary cells from patients with breast cancer also showed that rolipram could alter the enzymatic activities of MMP2 and MMP9, independent of the cAMP-PKA signaling pathway. These observations indicate the ability of rolipram to control breast cancer by regressing the functions of MMP2 and MMP9, thus having ‘off-targets’ other than PDE4 to have direct control over proteins that are involved in the advancement of metastasis.

## 1 Introduction

The complex and multistage cancer invasion and metastasis process is characterized by numerous genetic alterations. A crucial element in this progression is the breakdown of the extracellular matrix^1^. Matrix metalloproteinases (MMPs), a group of zinc-dependent endopeptidases with the ability to break down components of the extracellular matrix, play a significant role in facilitating cancer invasion and metastasis^2,3^. Increased concentrations of specific MMPs can be identified in tumor tissue or serum from individuals with advanced cancer^4,5^, therefore, their potential as prognostic markers in cancer has been extensively studied. Moreover, the expressions of MMP2 and MMP9, which are characterized as gelatinase, have been implicated in the progression and metastasis of breast cancer. Both MMPs are known to be differentially expressed in breast tumor tissues compared to adjacent normal tissues^6^. The overexpression of MMP2 and MMP9 was also associated with a poor overall survival in breast cancer patients, as well as higher histological grades, greater tumor size, and metastases^7^. MMP2^8^ and MMP9^9^ were also found to be responsible for the modulation of the immune response in the microenvironment of breast tumors. Therefore, several attempts have been made by researchers to functionally inhibit the activities of these MMPs to antagonize the progression and metastasis of breast cancer.

In a previous study, it was shown that the phosphodiesterase 4 (PDE4) inhibitor rolipram was able to alter the fate of the hedgehog signaling pathway in both the hormone-responsive breast cancer cell line MCF-7 and the triple-negative breast cancer cell line (TNBC) MDA-MB-231 in a cAMP-PKA-dependent manner^10^. Furthermore, rolipram was also observed to modulate the PTEN-PI3K-Akt axis in breast cancer cell lines^10^. Modulations of these pathways resulted in a successful down-regulation of MMP2 and MMP9 expression, along with other epithelial-to-mesenchymal transition (EMT) markers such as E-cadherin and vimentin in breast cancer cell lines. Concomitant retardations in the wound healing capabilities of both cell lines were also evident^11^. Since MMP2 and MMP9 expressions cannot justify their activities, the objective of this work has been to analyze the effect of rolipram on the enzymatic activities of these MMPs and to understand whether the mechanism of action of rolipram on MMP2 and MMP9 is always dependent on cAMP-PKA, as speculated in previous studies^11^.

The aforementioned study already predicted rolipram as an anti-breast cancer agent that can be exploited in combination with existing chemotherapeutic drugs to curb chemoresistance and tumor relapse. Hence, rolipram has been considered for drug repurposing to offer a shorter path in drug discovery for breast cancer. Although rolipram traditionally targets the phosphodiesterase 4 isoform, due to the intrinsic complexity of the molecular structure, drugs such as rolipram have the potential and practical interaction possibilities with several targets, which can be termed as “off-targets”, either in a particular signaling cascade or in different cascades. This idea of polypharmacology has changed the idea of drug design from having one target protein in a disease to having multiple target proteins for better efficacy. Hence, targeting druggable proteins might lead to the identification of novel off-target interactions, leading to a successful focus on drug repurposing. Additionally, the *in silico* prediction of ligand-protein interactions would reduce and refine the excessive budget of the laboratory processes significantly.

## 2 Results

### 2.1 Enzymatic activity of MMP2 and MMP9 in presence of rolipram

To assess the modulation of enzymatic activities of MMP2 and MMP9 in MCF-7 and MDA-MB-231 cell lines, gelatine zymography of the conditioned culture media was carried out. MCF-7 and MDA-MB-231 cells were treated with respective *IC*_50_ doses of 40 *μM* and 53 *μM* rolipram, which were determined in the previous study^11^. The data obtained from the study revealed that the activities of both MMP2 and MMP9 were significantly down-regulated by rolipram treatment. To observe whether the activities of these enzymes depend on cAMP-PKA, MCF-7 and MDA-MB-231 cells were treated with H89 (10 *μM*)^10^. It was evident from the zymography that H89 did not significantly inhibit the activity of both MMPs. However, both cell lines also showed significant inhibition of MMP2 and MMP9 activities after cotreatment with rolipram and H89, indicating a modulation of the MMP proteins independent of cAMP-PKA (Fig. 1 A).

**Figure 1.**
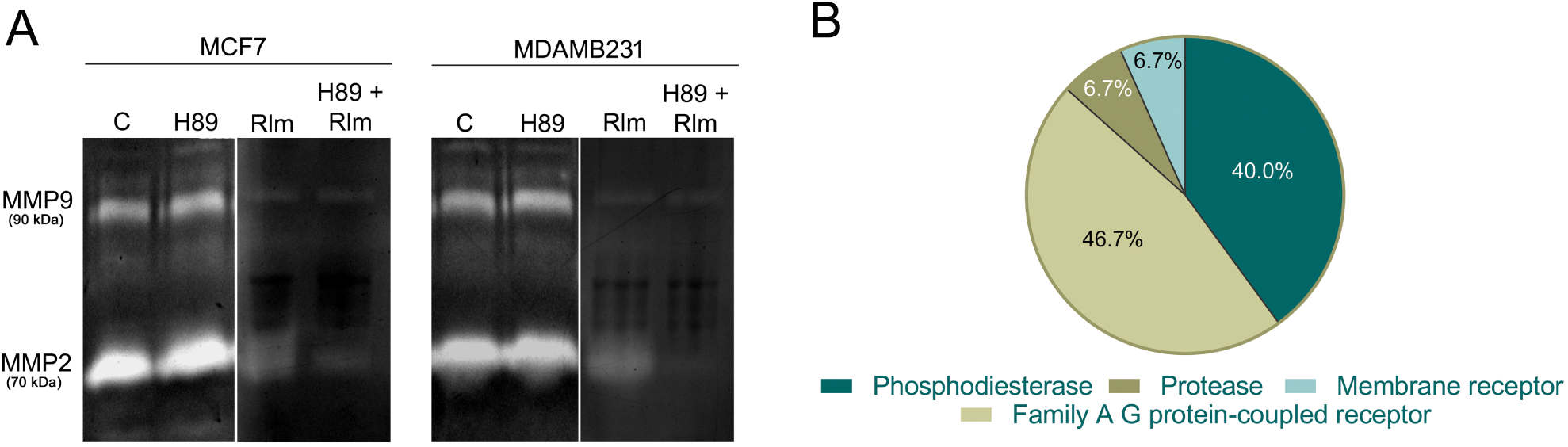
(A) Gelatin zymography of culture media conditioned with MCF-7 and MDA-MB-231 cells. The cells were treated with H89, rolipram (Rlm), and H89-rolipram for 24 hours. (B) Percentages of predicted probable protein targets (from *Homo sapiens*) of rolipram obtained from the SwissTargetPrediction server.

### 2.2 Determination of possible off-targets of rolipram

As rolipram showed an inhibitory effect on MMP2 and MMP9 even in the presence of H89, determining the mode of action of rolipram on MMP2 and MMP9 was of key importance. Therefore, it was crucial to identify possible off-targets of rolipram, which may mediate inhibitory effects even in the presence of H89. Furthermore, previous data obtained from gelatin zymography also raised the question of a possible direct interaction between rolipram and MMP proteins, bypassing the cAMP-PKA signaling pathway. The data revealed that 6.7% of the total probable targets of rolipram consist of proteases along with 40% comprising phosphodiesterases (Fig. 1 B, Table S1). This information led to the investigation of whether there are potential interactions between these MMPs and rolipram or not.

### 2.3 Prediction of MMP2 and rolipram interaction

Molecular docking studies were conducted to predict the potential interaction between rolipram and MMP2. In this study, 2-[(4- phenoxyphenyl)sulfonylmethyl]thiirane (SB3CT), a well-known inhibitor of MMP2 and MMP9 available on the market, was also analyzed as a positive control. The most favorable model obtained from Autodock was analyzed for each complex, MMP2 -rolipram and MMP2 - SB3CT (as a control). The best MMP2–rolipram model showed a binding affinity of *−*7.2 *kcal/mol*. Further analysis revealed two potential conventional hydrogen bonds between rolipram and threonine (THR) at the 56^*th*^ residue, as well as glutamine (GLN) at the 52^*nd*^ residue of MMP2. In addition, van der Waals interactions were observed between rolipram and MMP2 at the following sites: Tyrosine (TYR) in the 53^*rd*^ residue, phenylalanine (PHE) in the 57^*th*^ residue, methionine (MET) in the 97^*th*^ residue, proline (PRO) in the 43^*rd*^ residue and glycine (GLY) in the 189^*th*^ residue (Fig. 2 A). In the control simulation, the best model of the MMP2–SB3CT complex exhibited a binding affinity of *−*6.1 *kcal/mol*. This complex showed two conventional hydrogen bonds between SB3CT and GLN in the 323^*rd*^ residue, as well as leucine (LEU) at the 548^*rd*^ residue of MMP2, along with van der Waals interaction with serine (SER), THR and TYR residues (Fig. 2 B). Therefore, it can be inferred that rolipram may interact with MMP2 as effectively as SB3CT.

**Figure 2.**
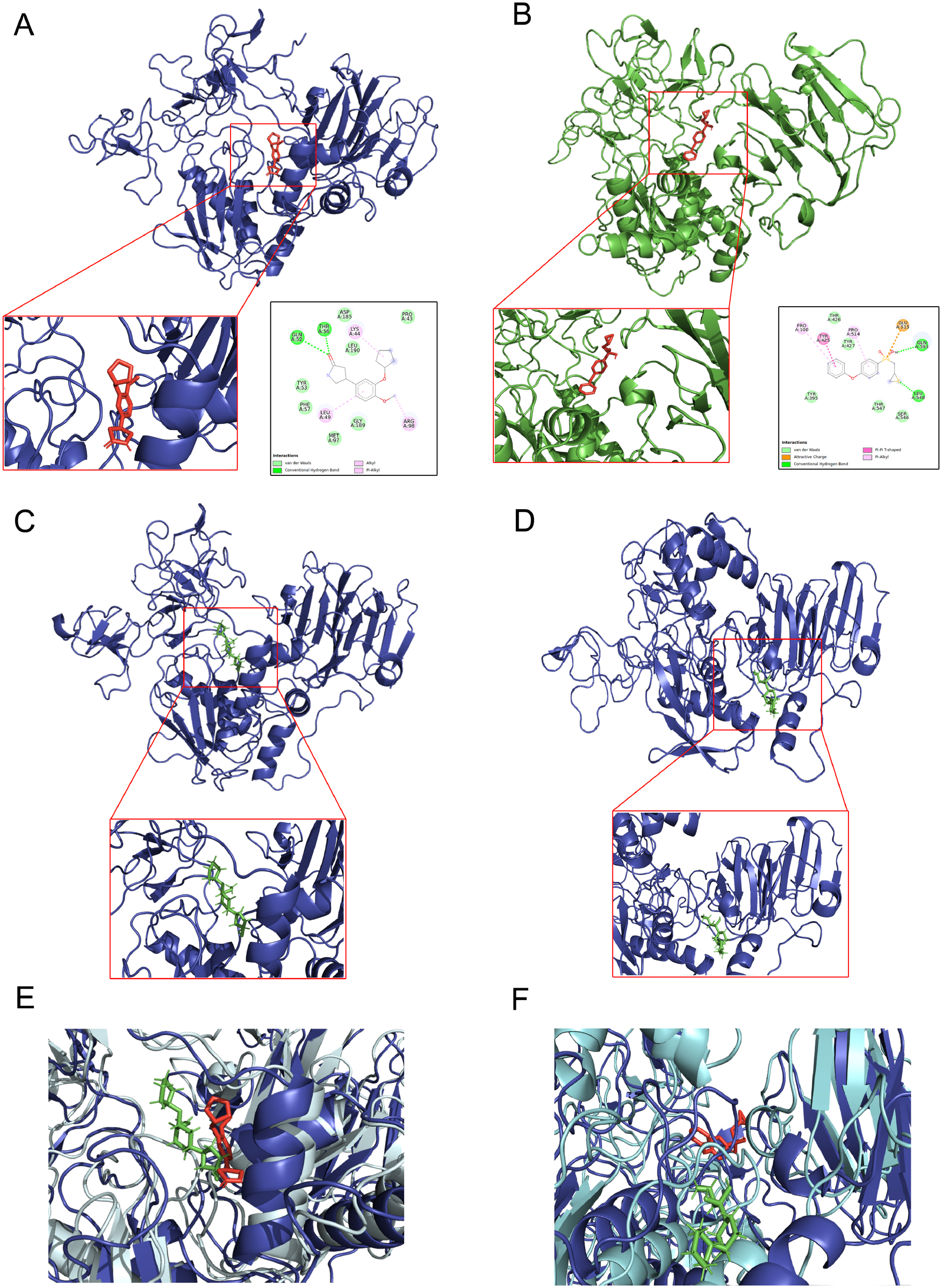
Top row: The most favorable docked complexes of MMP2-rolipram (panel A) and MMP2-SB3CT (panel B). The insets in the lower right display two-dimensional diagrams of the predicted interactions. Middle row: Snapshots of the MMP2-rolipram complex (panel C) and the MMP2-SB3CT complex (panel D), at the end of the 300 *ns* (nanosecond) MD simulation. Bottom row: Merger of the docked poses and the snapshot at the end of the simulation of the MMP2-rolipram complex (panel E) and MMP2-SB3CT (panel F).

### 2.4 Stability and integrity analysis of MMP2-rolipram and MMP2-SB3CT complexes

To further investigate the stability of the MMP2-rolipram and MMP2-SB3CT complexes, molecular dynamics (MD) simulations were performed on the best model complexes obtained from molecular coupling. Furthermore, a simulation of the free MMP2 protein was conducted without ligands to study its dynamics. Simulations were performed for 300 *nanoseconds* (*ns*) to examine the stability and dynamic behavior of these complexes over time.

During simulation, the position of the rolipram molecule in the MMP2-rolipram complex remained unchanged (Fig. 2C And 2E). However, it was observed that the SB3CT molecule in the MMP2-SB3CT complex shifts to the adjacent MMP2 binding pocket at the end of the simulation (Figs. 2 D and 2 F). Analysis of the root-mean square deviation (RMSD) showed that the free MMP2 protein achieved stability around 10 *ns*, with an RMSD of 0.4 *nm* from its initial configuration. On the other hand, the MMP2-rolipram complex did not stabilize within the first 100 *ns*, and its RMSD fluctuated between 0.4 and 0.6 *nm*. In contrast, the MMP2-SB3CT complex reached stability within 10 *ns*, with an RMSD of 0.8 *nm* from its initial configuration. Eventually, the MMP2-rolipram complex stabilized around 100 *ns* with an RMSD of approximately 0.4 *nm* (Fig. 3A). Analysis of the radius of gyration (Rg) showed that the MMP2 protein retained its compactness in the presence of both ligands, as well as in its free form. The MMP2-rolipram complex maintained its initial compactness for the first 40 *ns* of the simulation with fluctuations of less than 0.05 *nm* from a locally time-averaged value of 2.85 *nm*. The Rg then fell below this average between 40 and 100 *ns*, before rising and stabilizing at 2.8 *nm* until the end of the simulation.

**Figure 3.**
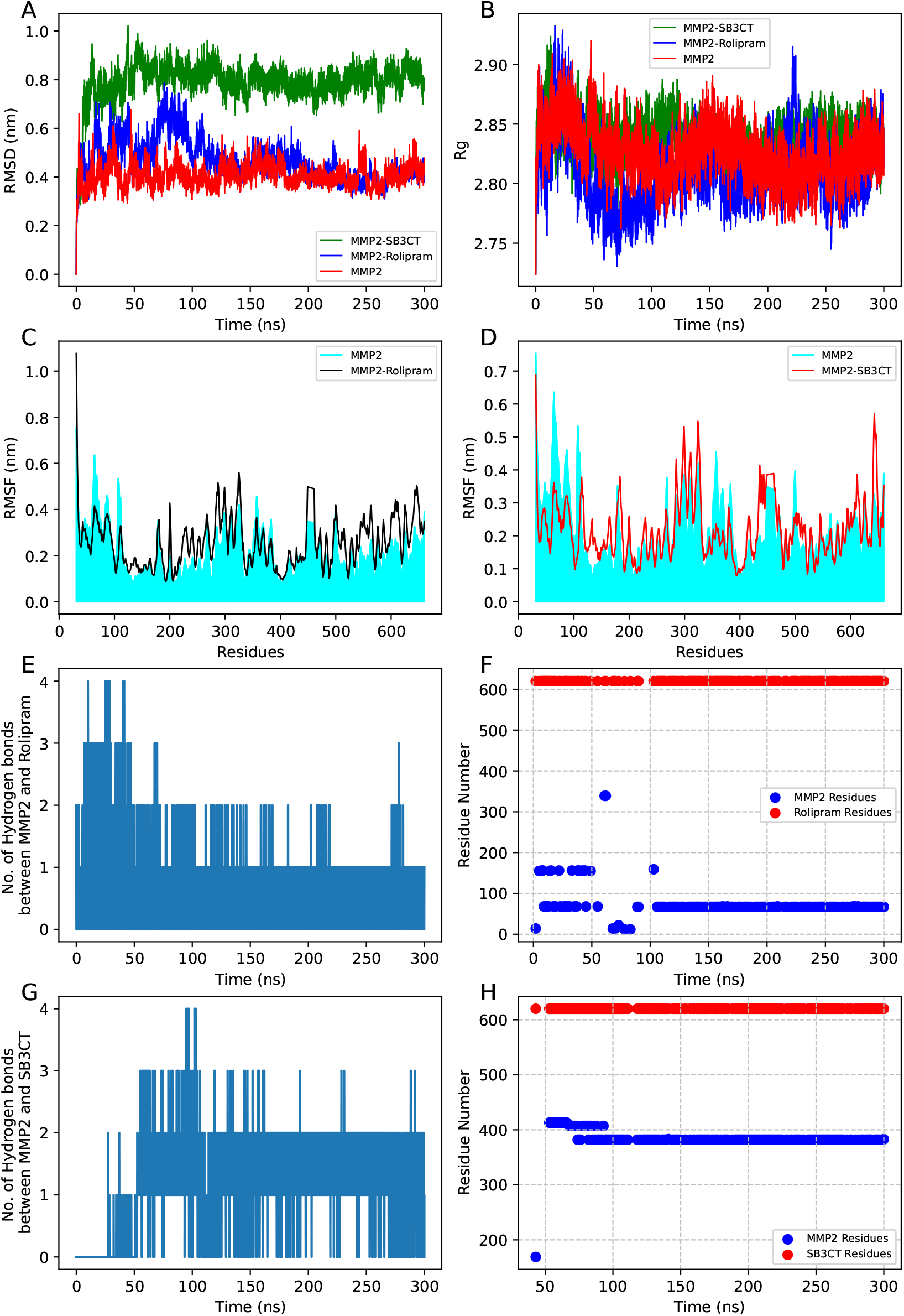
First row at the top: plots of root-mean square deviation (RMSD) versus time of the free MMP2, MMP2-rolipram complex, and MMP2-SB3CT complex (panel A), as well as plots of the radius of gyration (Rg) of the same (panel B). Second row: plots of root mean square fluctuation (RMSF) of the MMP2-rolipram complex (panel C) and the MMP2-SB3CT complex (panel D), both with respect to free MMP2, for 300 ns of simulation time each. Third row: Time plots of the number of hydrogen bonds between MMP2 and rolipram that were detected by GROMACS (panel E), as well as the identification numbers of the protein residues that MDAnalysis revealed to have participated in hydrogen bonds (panel F). Last row at the bottom: Similar plots for the MMP2-SB3CT complex, the number of hydrogen bonds (panel G) and the residue numbers (panel H).

In the control simulation, the MMP2-SB3CT complex remained stable throughout, with minimal fluctuations, at an Rg of 2.85 *nm*. Interestingly, the free MMP2 simulation exhibited even greater stability in Rg, with minimal fluctuations ranging from 2.8 to 2.85 *nm* within the same simulation time (Fig. 3B). Further analysis of root mean square fluctuation (RMSF) of specific amino acid residues in these complexes, compared to the free MMP2 simulation, provided insights into possible structural changes of the MMP2 protein due to its interactions with ligands. Significant fluctuations of approximately 0.2 *nm* were observed in a specific range of amino acid residues (from 10 to 100) in MMP2. The SB3CT ligand caused significant fluctuations of 0.2 *nm* in a different range of amino acid residues (from 330 to 380) in MMP2. Furthermore, both ligands induced fluctuations ranging from 0.1 to 0.2 *nm* in residues 420 to 600 of MMP2 (Fig. 3C, 3D and S7A).

### 2.5 Identification and analysis of stability of hydrogen bonds involving rolipram and SB3CT with MMP2

Analysis of hydrogen bonds formed between MMP2 and rolipram during the simulation indicated that hydrogen bonds were present from the beginning. Between 10 *−* 40 *ns*, the GROMACS `hbond` module identified at least two stable hydrogen bonds between MMP2 and rolipram. After 40 *ns*, at least one stable hydrogen bond was identified and was shown to remain throughout the rest of the simulation (Fig. 3E). Using the MDAnalysis tool in the trajectory data files, it was discovered that the first two hydrogen bonds were between the oxygen atom of the rolipram molecule and the 68^*th*^ ARG residue of MMP2 and the nitrogen atom of rolipram with the 156^*th*^ GLY residue of MMP2 (Fig. 3F). The subsequent stable hydrogen bond was between the nitrogen atom of rolipram and the oxygen atom of the 67^*th*^ residue of MET MMP2, starting from 100 *ns*. The angle between the hydrogen, donor and acceptor atoms of the 67^*th*^ MET, 68^*th*^ ARG and 156^*th*^ GLY residues of MMP2 and rolipram remained within the 30° cut-off (determined according to the Luzar and Chandler criterion^12,13^) throughout the simulation (Fig. S8 A, S8 B and S8 C). However, the distance deviation data showed that the distance between the donor and acceptor atoms of the 156^*th*^ GLY residue fell below the cutoff distance of 0.35 *nm* (according to Luzar Chandler) only from 0 to 50 *ns*. Similarly, the distance between the acceptor atom of the rolipram molecule and the donor atom of the 68^*th*^ ARG residue of MMP2 was also within the cutoff distance at the same times. Furthermore, the distance between the oxygen atom from the 67^*th*^ MET residue of MMP2 and the nitrogen atom of rolipram was within the cutoff range during simulation times from 100 to 300 *ns* (Fig. S8 D, S8 E and S8 F).

Similarly, the analysis of hydrogen bonds between MMP2 and SB3CT in the control simulation revealed that there were at least two stable hydrogen bonds between MMP2 and SB3CT that began at 50 *ns* (Fig. 3G). The MDAnalysis tool revealed that two atoms of the 382^*nd*^ GLU residue of MMP2 are involved in stable hydrogen bonds with SB3CT (Fig. 3H). The angle between the hydrogen, donor, and acceptor atoms of these residues remained within the cut-off point after 50 *ns* (Figs. S9A And S9B). The distance between these atoms was also within the cut-off during this time (Fig. S9C And S9D).

### 2.6 Prediction of interaction between MMP9 and rolipram

Docking studies between rolipram and MMP9 were initially carried out following a methodology similar to that for MMP2 as described in Subsection 2.3. The analysis indicated that the most favorable MMP9-rolipram complex had a binding affinity of *−*7 *kcal/mol*. Examination of the MMP9-rolipram complex model for protein-ligand interaction revealed the presence of two potential conventional hydrogen bonds. A hydrogen bond was identified between rolipram and the 51^*st*^ ARG residue of MMP9, and the other with the 96^*th*^ THR residue (see Fig. 4A). In the control docking, the best MMP9-SB3CT complex displayed a binding affinity of *−*6.2 *kcal/mol*, although no conventional hydrogen bond was predicted between the two entities. However, several non-bonding van der Waals interactions were observed, involving the SB3CT molecule and various MMP9 residues, including LEU in the 44^*th*^ residue, TYR in the 48^*th*^ and 52^*nd*^ residues and ARG in the 51^*st*^ residue (see Fig. 4B).

**Figure 4.**
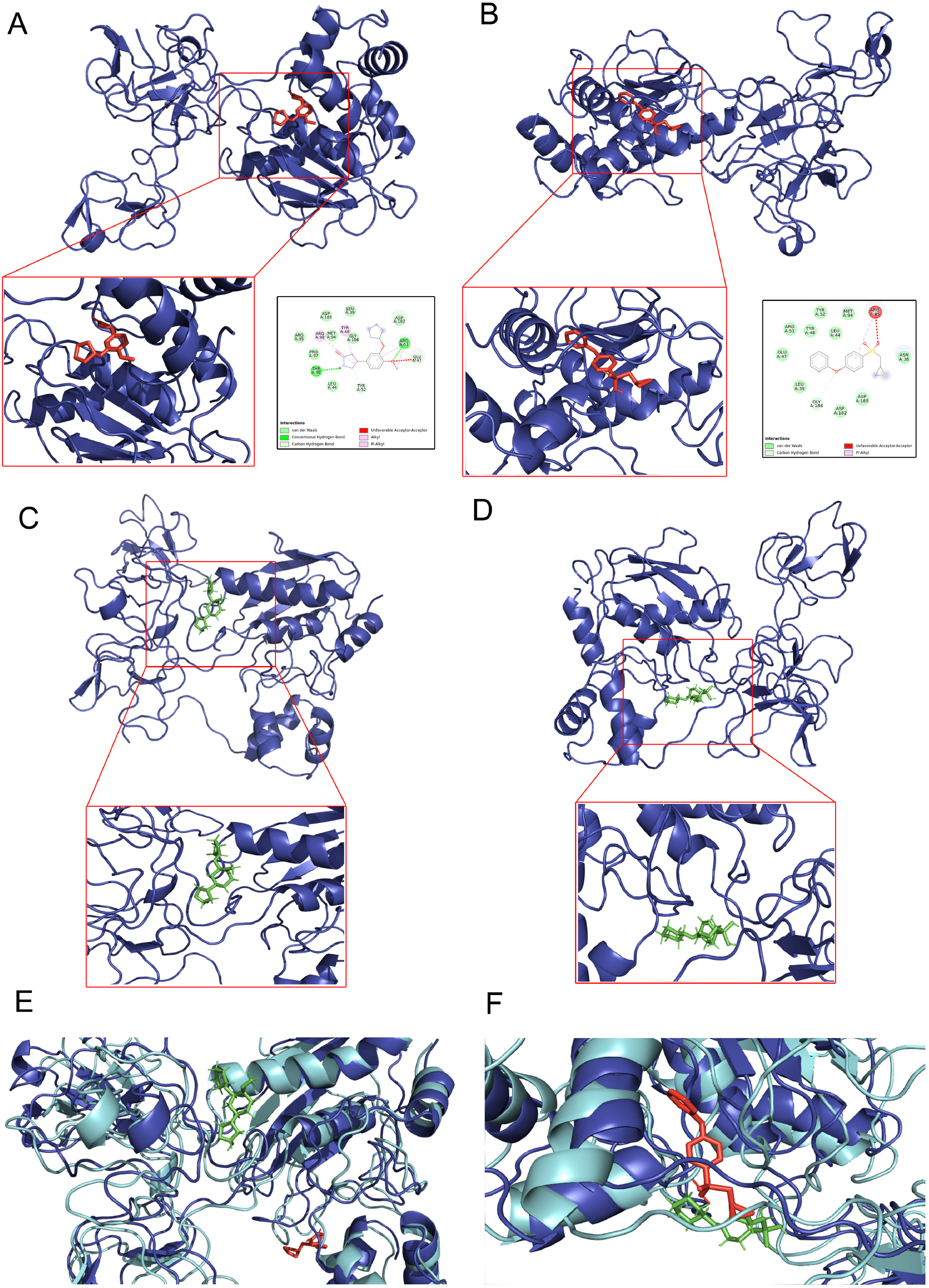
Top row: The most favorable docked complexes of MMP9-rolipram (panel A) and MMP9-SB3CT (panel B). The insets in the lower right display two-dimensional diagrams of the predicted interactions. Middle row: Snapshots of the MMP9-rolipram complex (panel C) and the MMP9-SB3CT complex (panel D), at the end of the 300 *ns* MD simulation.

### 2.7 Stability and integrity analysis of MMP9-rolipram and MMP9-SB3CT complexes

The stability of MMP9 interactions with rolipram and the SB3CT control ligand was examined by MD simulations *in silico*, using parameters similar to those of MMP2 and ligands detailed in Section 2.4. The MMP9-rolipram complex that exhibited the most favorable docking was selected as the starting point of the simulation. Comparison of the molecular pose after 300 *ns* simulation with the initial configuration showed that the rolipram molecule had moved to an adjacent binding site in MMP9 before reaching stability (Figs. 4 C and 4 E). In the control simulation, the SB3CT molecule was observed to alter its position within the same binding site and stabilize within the same time frame (Fig. 4D And 4F). The simulation involving free MMP9 appeared to stabilize at an RMSD of 0.4 *nm* after 30 *ns*, but showed more noticeable deviations after 80 *ns*. However, RMSD values later decreased, returning to earlier levels after 100 *ns*. In the simulation of the MMP9-rolipram complex, the protein stabilized (within 10 *ns*) at an RMSD of 0.4 *nm*, while the MMP9-SB3CT complex initially stabilized at an RMSD of 0.3 *nm* during the same period, then increased to 0.4 *nm* after 80 *ns*, remaining stable for the rest of the simulation (Fig. 5A).

**Figure 5.**
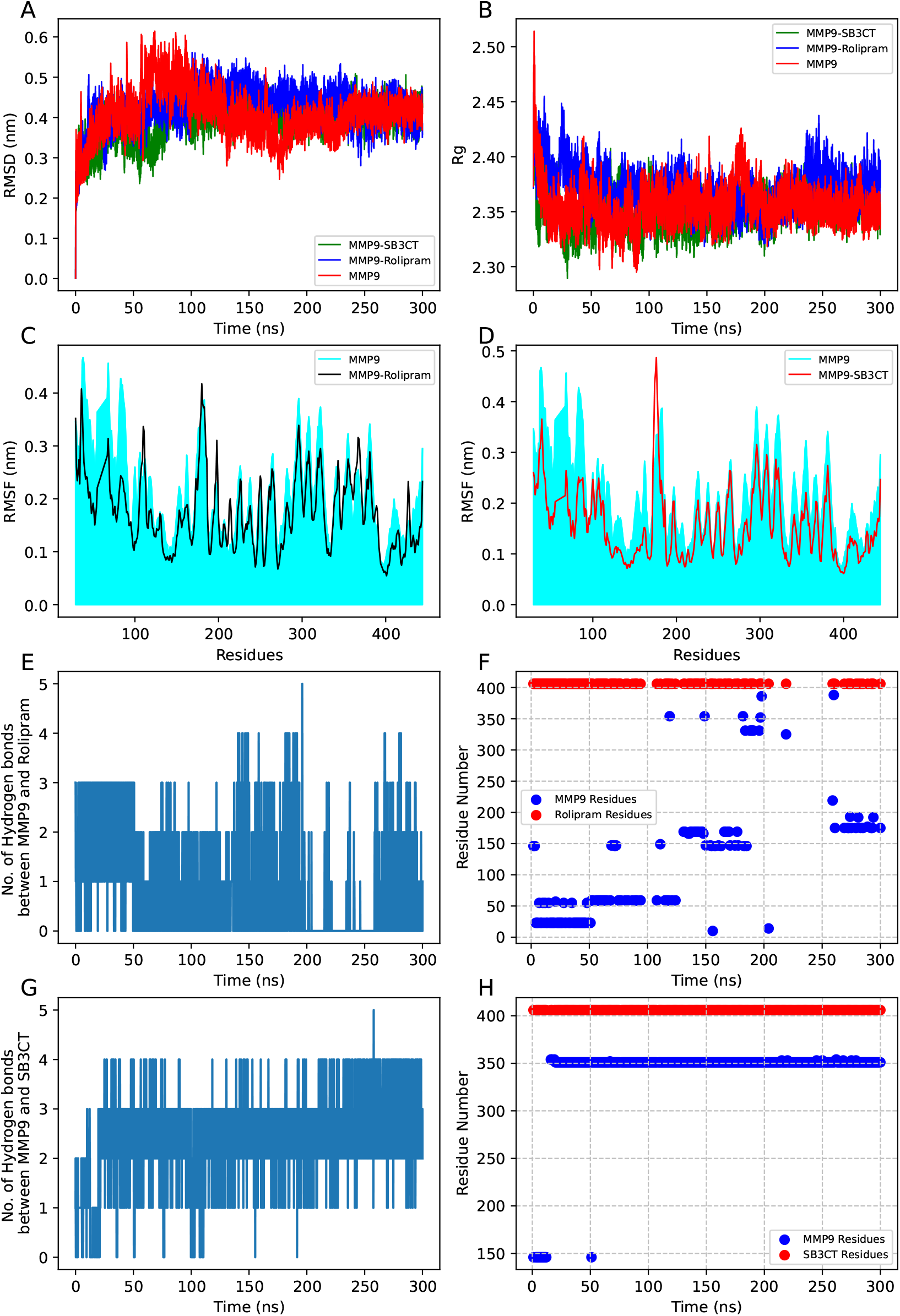
First row at the top: RMSD plots versus time of the free MMP9, MMP9-rolipram complex, and MMP9-SB3CT complex (panel A), as well as plots of the radius of gyration (Rg) of the same (panel B). Second row: RMSF plots of the MMP9 complex (panel C) and the MMP9-SB3CT complex (panel D), both with respect to free MMP9, for 300 *ns* of simulation time each. Third row: Time plots of the number of hydrogen bonds between MMP9 and rolipram that were detected by GROMACS (panel E), as well as the identification numbers of the protein residues that MDAnalysis revealed to have participated in hydrogen bonds (panel F). Last row at the bottom: Similar plots for the number of hydrogen bonds identified by GROMACS with respect to time for the MMP9-SB3CT complex (panel G) and the protein residues identified by MDAnalysis (panel H).

Analysis of the Rg plots of the complexes and free MMP9 showed that the Rg values of free MMP9 and MMP9 in complex with SB3CT stabilized at 2.35 *nm*, with a slight fluctuation of 0.05 *nm*, while the Rg of MMP9 in complex with rolipram stabilized nearby at around 2.38 *nm* (Fig. 5B). Examination of the residue-specific RMSF indicated that rolipram and SB3CT induced notable fluctuations (compared to free MMP9) of approximately 0.1 to 0.2 *nm* in MMP9 amino acid residues spanning 20 to 100 and the range of 280 to 320. Furthermore, minor fluctuations in amino acid residues were observed beyond the 400^*th*^ residue of MMP9 in both complexes. The SB3CT molecule also caused fluctuations in MMP9 amino acid residues ranging from 320 to 380, while these residues were not affected in the presence of rolipram (Fig. 5C, 5D and Fig. S7B).

### 2.8 Identification and examination of the stability of hydrogen bonds between rolipram and SB3CT with MMP9

Analysis of hydrogen bonds formed between MMP9 and rolipram showed that there were at least three stable hydrogen bonds between these two up to 50 *ns* of simulation time. Subsequently, there was at least one stable hydrogen bond between MMP9 and rolipram up to 200 *ns* (Fig. 5E). Another stable hydrogen bond was observed after 260 *ns*.

The MDAnalysis tool detected a minimum of five residues in MMP9 that participated in hydrogen-bonding interactions with rolipram. Among these, the 23^*rd*^ ARG residue was observed to form bonds with the oxygen atom of rolipram for up to 50 *ns*. Another residue, 59^*th*^ ARG of MMP9, exhibited a stable bond with an oxygen atom of rolipram from 50 to 130 *ns*. Furthermore, the 169^*th*^ GLU and 146^*th*^ ASP residues of MMP9 were identified to be involved in hydrogen bonds with the oxygen and nitrogen atoms of rolipram, respectively, from 130 *ns* to 190 *ns*. Subsequently, at 250 *ns*, the 175^*th*^ LYS residue was found to bond to the oxygen atom of rolipram (Fig. 5F). Detailed angular deviation plots of the angles between the hydrogen, donor, and acceptor atoms of these residues showed minimal variation throughout the simulation (Fig. S10A-E). However, the distance deviation plots indicated that the distance between the donor and acceptor atoms in the 23^*rd*^ ARG residue of MMP9 and rolipram remained within the cutoff distance only during the initial 50 *ns* before dissociation.

Between 50 and 125 *ns* of the simulation time, the distance measurements of the hydrogen bond-forming oxygen atom in rolipram and the 59^*th*^ ARG residue of MMP9 were within the Luzar-Chandler cutoff criteria, after which the bonds were broken. Similarly, the distance related to hydrogen bond formation between rolipram and 146^*th*^ ASP, as well as the 169^*th*^ GLU residue of MMP9, remained within the cut-off thresholds during the time intervals 150 *−* 190 *ns* and 125 *−* 170 *ns*, respectively.

During simulation, a new hydrogen bond emerged as the distances between rolipram and the 175^*th*^ LYS residue of MMP9 decreased below the cutoff value from 260 *ns* (see Fig. S10 FJ).

In the control simulation, GROMACS identified a minimum of three stable hydrogen bonds between MMP9 and SB3CT starting at 20 *ns* (Fig. 5G). According to MDAnalysis, the 351^*st*^ ASP residue of MMP9 was involved in a hydrogen bond with SB3CT from 20 *ns* (Fig. 5H). MDAnalysis also detected that two neighboring oxygen atoms of the 351^*st*^ ASP residue of MMP9 were simultaneously creating hydrogen bonds with the oxygen atom of SB3CT. The angles involving the hydrogen, donor and acceptor atoms of the two hydrogen bonds were consistently within the predefined range throughout the simulation period, while the distance between the donor and acceptor atoms dropped below the threshold at 20 *ns* and remained within the limit for the remainder of the simulation period (Fig. S11).

### 2.9 Rolipram inhibited gelatinase activity of MMP2 and MMP9 in primary cells of breast cancer patients

Following the simulations of the computational models, the interaction of rolipram with MMP2 and MMP9 was further substantiated by gelatin zymography of culture media conditioned with primary cells of tumors of patients with breast cancer. The conditioned media of the primary tissue cultures were collected and incubated with rolipram at concentrations of 20 and 40 *μM*, each for 3 hours. Since the culture medium was already devoid of cells prior to rolipram treatment, the application of the PKA inhibitor H89 was not necessary to validate the activity of rolipram. Zymography showed that the enzymatic activities of MMP2 or MMP9 were not exerted on gelatin in the presence of rolipram, while both MMPs were able to exert their enzymatic activities in the absence of rolipram (Fig. 6).

**Figure 6.**
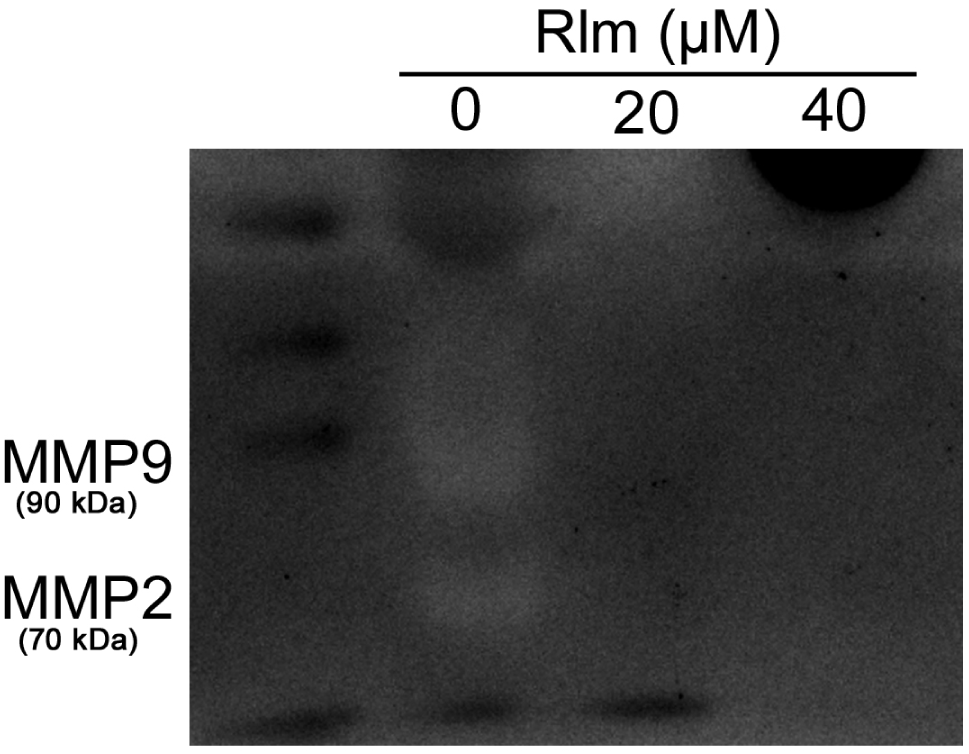
Gelatin zymography of culture media conditioned with primary cells from human breast tumor tissue. The conditioned media were incubated with 20 and 40 *μM* concentrations of rolipram for 3 hours before zymography

## 3 Discussion

A previous investigation (available at^11^) demonstrated that rolipram negatively affected cell migration and scratch wound healing through hedgehog signaling dependent on cAMP-PKA and cAMP-PKA/PI3K-Akt signaling pathways. Rolipram showed modulations in intracellular levels of MMP2 and MMP9, which are known to be associated with cell migration and metastatic potential of cells. However, cAMP-PKA-dependent modulations of MMP2 / 9 expression levels could not conclusively demonstrate the enzyme activities of these proteins. To further explore whether the regulation of MMP2 and MMP9 activities was solely cAMP-PKA-dependent (as apparently thought from studying the expression profiles of MMP2 and MMP9) gelatin zymography was performed with rolipram after co-administration of H89, a potent PKA inhibitor. The results of the zymography analysis showed that rolipram effectively suppressed the gelatinase activity of MMP2 and MMP9 even in the presence of H89, challenging the previous assumption that such modulations were cAMP-PKA-dependent. This outcome suggested the possible potential of rolipram to influence MMPs by circumventing the cAMP-PKA-mediated signaling cascade. Rolipram was screened for potential off-target effects, revealing that some specific proteases, such as MMPs, were prominent in the screening results, along with its primary targets, PDEs.

Based on this prediction, possible interactions between rolipram and MMP proteins *in silico* were investigated. Molecular docking studies demonstrated significant binding affinities of rolipram with MMP2 and MMP9, similar to the control ligand, SB3CT, which is known to be a potent inhibitor of both MMP2 and MMP9. To further support the hypothesis that rolipram interacts with MMP2 and MMP9, MD simulations of the best-docked complexes were performed. The simulation results offered valuable information on the likely interaction of rolipram with MMP2 and MMP9. Both proteins, in their free state or in the presence of rolipram, remained stable and compact throughout the simulation period, mirroring the control simulation with SB3CT. During simulations, rolipram was observed to shift its initial docked position with MMP2 and establish a stable hydrogen bond. In the case of MMP9, rolipram formed multiple stable hydrogen bonds at various time points after moving away from the initial docked configuration. As expected, SB3CT also showed stable hydrogen bond formation with both proteins throughout the simulation, along with noticeable deviations from the initial docked conformations.

Therefore, the results of the MD simulations of these complexes provided improved insights into the functioning of rolipram, a phosphodiesterase 4 inhibitor. Furthermore, when rolipram was added to the conditioned medium of the primary cells obtained from human breast cancer patients, it inhibited the enzymatic activities of MMP2 and MMP9. The conditioned medium was incubated with 20 and 40 *μM* concentrations of rolipram. Because this medium was cell-free, it was evident from the result that rolipram’s effect on the activities of these MMPs was independent of the cAMP-PKA signaling pathway affecting cancer cells in the primary culture. Thus, the findings of this study depict a possible off-target of the phosphodiesterase 4 inhibitor rolipram, suggesting its potential as a promising option for breast cancer therapy targeting proteins that are potent markers of metastasis.

## 4 Methods

### 4.1 Ethics Statement

No animals were used in this study. Human breast cancer tissue samples were collected from the All India Institute of Medical Sciences (AIIMS) Kalyani, Kalyani, Nadia, West Bengal, India, in strict compliance with the guidelines of the Institutional Human Ethics Committee of AIIMS Kalyani.

### 4.2 Materials

The estrogen and progesterone receptor-positive human breast cancer cell line MCF-7 and the triple-negative human breast cancer cell line MDA-MB-231 were purchased from the National Center for Cell Sciences (NCCS), Pune, India. The PDE4 inhibitor, Rolipram, purchased from Sigma-Aldrich (USA, Cat # R6520) and a stock solution of 10 *mM* was prepared by dissolving it in ethanol. PKA inhibitor H89 was also purchased from Sigma-Aldrich (USA, Cat # B1427) and dissolved in DMSO to prepare a stock solution of 1 *mM*. Dulbecco’s modified eagle medium (DMEM, Cat # 11885084), fetal bovine serum (FBS, United States, Cat # 16140071), penicillin-streptomycin (Cat # 15140122) were purchased from Gibco™ (USA).

### 4.3 Cell culture and treatment

Human breast cancer cell lines were maintained in a culture medium consisting of DMEM, supplemented with 10% FBS and 100 *U/ml* penicillin-streptomycin. The cells were kept in a humidified atmosphere 5% CO_2_ at 37°*C*. The cells were checked for the presence of any contaminants before any experiment and allowed to attain at least 70% confluence^14^. The MCF-7 and MDA-MB-231 cells were treated with 40 *μM* and 53 *μM* concentrations of rolipram, respectively.^11^ for 24 hours. 10 *μM*^10^ concentration of H89 was also used to treat these cell lines separately and in combination with previously mentioned doses of rolipram.

### 4.4 Primary tissue culture from human breast tumor tissue

The human breast tumor tissue samples were obtained from AIIMS Kalyani after receiving informed consent from patients according to the AIIMS Institutional Human Ethical Committee guidelines. The tissues were obtained from Tru-cut biopsy of the patients which were authenticated to be breast cancer by trained pathologists. The samples were washed with antibiotics (100 U/ml penicillin, 100 *μ*g/ml streptomycin) and incubated with the collagenase-hyaluronidase mixture for 16-18 hours at 37°*C* before being maintained in a culture medium, DMEM/F-12, supplemented with 10% FBS, 5*μg/ml* insulin and 0.5*μg/ml* hydrocortisone. For zymography, the cells were then maintained in a serum-free culture medium for 12 hours before collecting the culture medium, which was then equally divided into three microcentrifuge tubes. Two of these samples were incubated with 20 *μM* and 40 *μM* concentrations of rolipram for 3 hours at 37°*C* prior to the gelatin zymography study, while no incubation was carried out in the other sample.

### 4.5 Gelatin zymography

Gelatin zymography in MCF7 and MDA-MB-231 involved cell culture media conditioned by the human breast cancer cell lines treated with rolipram, H89, and co-treatment of H89 and rolipram for a 24 hour period. Gelatin zymography was also performed with culture medium conditioned with primary cells from tumor tissues from human breast cancer patients, incubated with two different doses of rolipram after collection of the cell-free medium after 3 h of incubation. They were first resolved using a nonreducing 10% polyacrylamide gel, containing 1 *mg/ml* gelatin. The gels were washed twice with 2.5% Triton™ X*−*100 (30 *minutes* each) and incubated with incubation buffer for 24 *hours* at 37°*C*. The incubation buffer, also known as the calcium assay buffer, is composed of 40 *mM* Tris-HCl (*pH* 7.4), 0.2 *M* NaCl, 10 *mM* CaCl_2_ and NaN_3_. Following incubation, the gels were stained with 0.1% Coomassie blue followed by destaining. Gel images were obtained from the Bio-Rad ChemiDoc XRS + System, using Bio-Rad Image Lab software, and zymographic band quantification was performed using the NIH ImageJ software^15,16^.

### 4.6 Drug target prediction

The canonical Simplified Molecular Input Line Entry System (SMILES) format of rolipram was obtained from PubChem^17^ and provided to the SwissTargetPrediction server^18^ in order to predict probable protein targets of rolipram.

### 4.7 Molecular docking

The structure of the MMP2 and MMP9 proteins were obtained from the Protein Data Bank (PDB) files with the PDB ids 1*ck*7 and 1*l*6 *j*, respectively, from the website www.rcsb.org. The 3D coordinates of the ligands, rolipram and 2-[(4-phenoxyphenyl)sulfonylmethyl]thiirane (SB3CT) were obtained from PubChem^17,19^ in the format of a structured data file (SDF). These coordinates were then converted to the PDB format using Open Babel^20^. To perform the computational molecular docking of MMP2 and MMP9 with rolipram and SB3CT, Autodock4 was used in AMDock (Version 1.6.x for Linux)^21^. The structures of the MMP2 and MMP9 protein, as well as the rolipram and SB3CT ligands, were provided as separate PDB files to AMDock. The input was prepared by loading the force field data using the CHARMM force field model, hydrogen topology data, and updated disulfide bridge data. No repairs were needed during this process. After input preparation, the search space for the minimum-energy configuration was defined using the autoligand function. Then, the Autodock software was instructed to perform independent docking simulations for every site predicted by autoligand. Finally, the output, which included the final poses of the protein-ligand complexes and their binding affinity in kilocalories per mole (*kcal/mol*), was recorded in the PDB files. To analyze the results, the protein-ligand interaction surfaces were visualized using Biovia Discovery Studio software.

### 4.8 Molecular dynamic simulation

Molecular dynamics (MD) simulations were performed using GROMACS (version 2022.4)^22–24^ to study the behavior of protein-ligand complexes. The simulations were carried out on a Terascale High-Performance Computing Cluster at the S.N. Bose Innovation Centre, University of Kalyani, Kalyani, India. GROMACS is a versatile molecular dynamics simulation package that can simulate complex biomolecules in virtual environments that mimic laboratory conditions. The molecular coordinates of the MMP2 and MMP9 proteins were converted to the appropriate simulation formats using internal GROMACS tools, as well as the CHARMM force field^25^. The ligands, rolipram and SB3CT, were generated using the SwissParam server^18^. The molecular coordinates of the proteins and ligands, along with solvent water molecules (whose microscopic chemistry was treated using the flexible simple point-charge water model) and charge-neutralizing ions, were combined to create a configuration file for each protein-ligand complex. The configuration file was combined with the force field parameters and external bath couplings (to achieve thermal equilibrium under laboratory conditions) to form a topology file that was fed to the GROMACS MD trajectory builder. Subsequently, GROMACS was instructed to randomly draw atom velocities from a Maxwell-Boltzmann distribution at room temperature and simulate the ensuing dynamics using the leapfrog numerical integration technique.

Thus, GROMACS was able to successfully generate trajectory files that captured the temporal dynamics of the atoms. These trajectory files were then analyzed using the GROMACS internal tools and the MDAnalysis Python module^26^. The stability and integrity of each protein-ligand complex were individually evaluated by calculating the root-mean square deviation (RMSD) of the protein backbone from its initial configuration and the radius of gyration (Rg) of the protein backbone throughout the simulation^27^. To assess structural fluctuations resulting from protein-ligand interactions, root-mean square fluctuations (RMSF) of the amino acid residues of the proteins were calculated. Higher RMSF values indicate regions with greater structural mobility. The stability and characteristics of the protein-ligand interaction were examined by analyzing the formation of hydrogen bonds. The number of hydrogen bonds formed during the simulation was determined using the shell command `gmx hbond`, and the MDAnalysis python module (Script Ref:^28^) was used to identify the acceptor and donor atoms involved in hydrogen bond formation. The distance and angle between the residues involved in hydrogen bond formation were analyzed over time using shell commands, `gmx distance` and `gmx angle`, respectively.

## Supporting information

LaTeX Source and figs

## Acknowledgements

The authors thank the Science & Engineering Board, Government of India [Grants SUR/2022/001275 and CRG/2018/004002], Department of Biotechnology, Government of West Bengal, India [No. 248 (Sanc)/BT (Estt)/RD-27/2016] for funding this study. The authors also thank the HPC Facility of the University of Kalyani, the DST-FIST and UGC-SAP programs of the Department of Zoology of the University of Kalyani, and the DST-PURSE-sponsored instrument facilities of the University of Kalyani for providing infrastructure facilities and the Personal Research Grant of the University of Kalyani.

## Author contributions statement

A. Bagchi performed all the experiments and simulations. A. Roy helped with HPC management and the preparation of the simulation scripts. A. Biswas & A.Roy obtained funding, conceptualized the experiments, designed them, interpreted the data, and prepared the manuscript. A. Halder provided the tru-cut biopsy sample of the breast cancer patient from which the primary cell culture was performed. All authors approve of this version of the manuscript as final.

## Additional information

### Data availability statement

All the data that support the findings of the study are available from the corresponding authors upon request.

## Supplementary Table and Figures

**Table 1.**
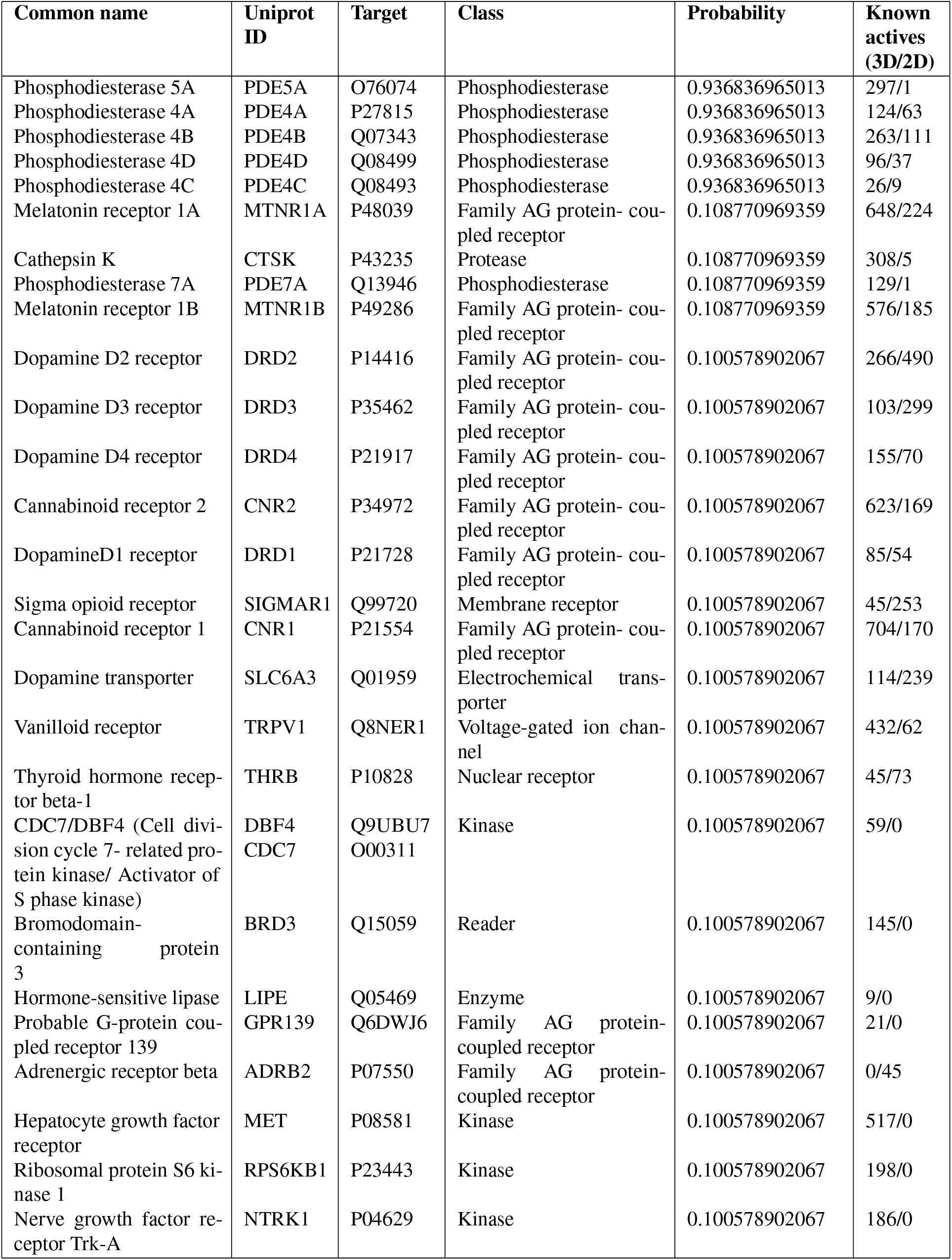
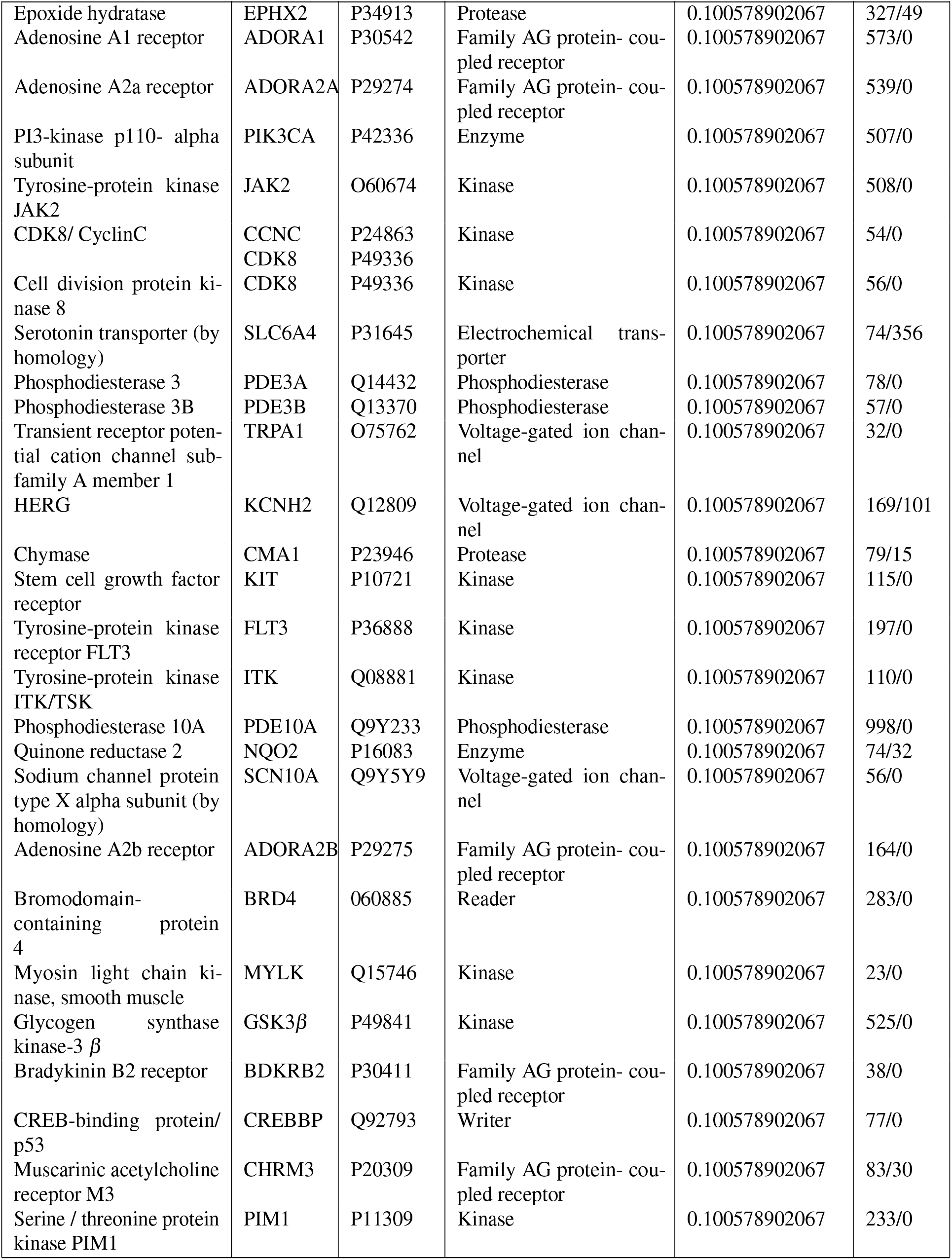
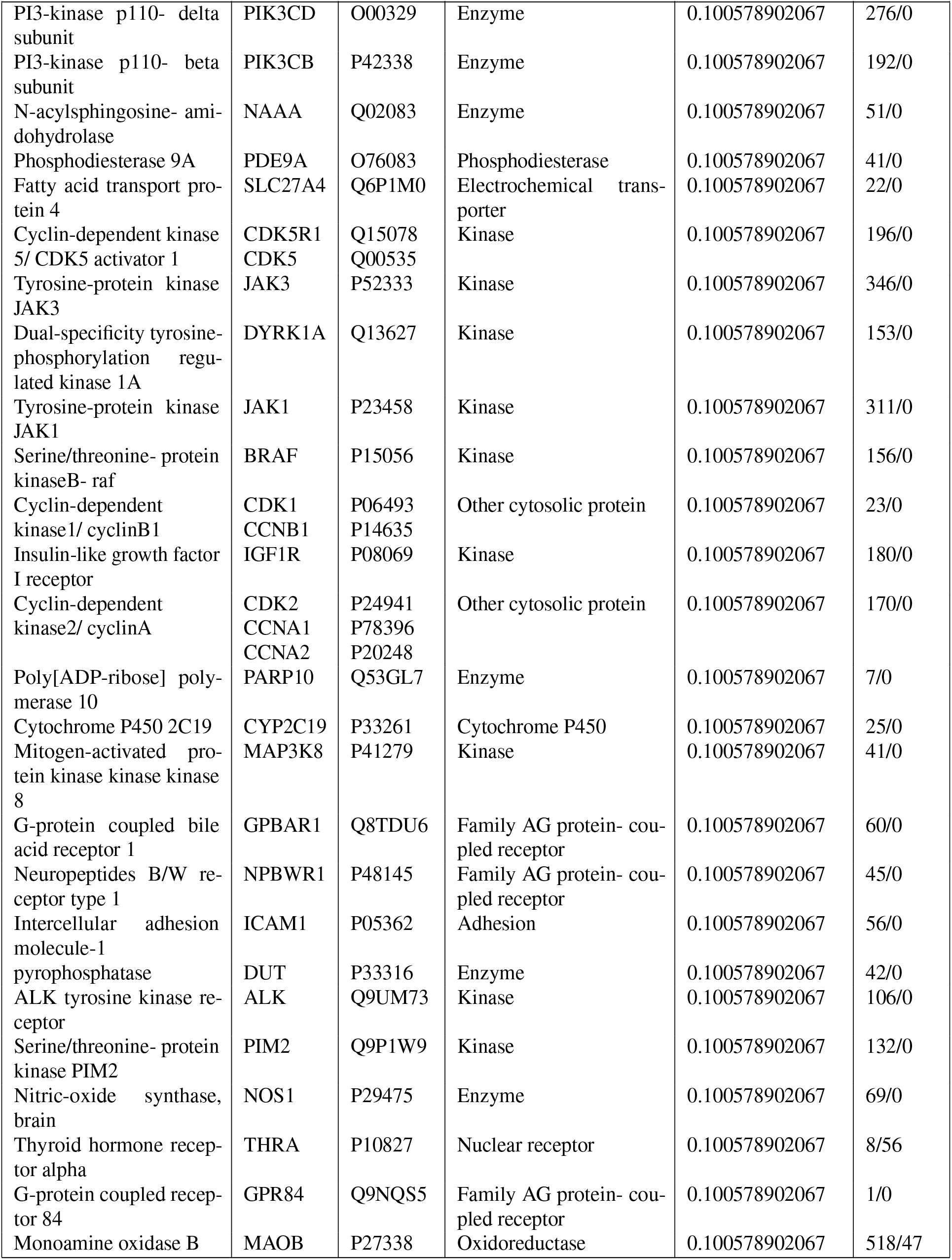
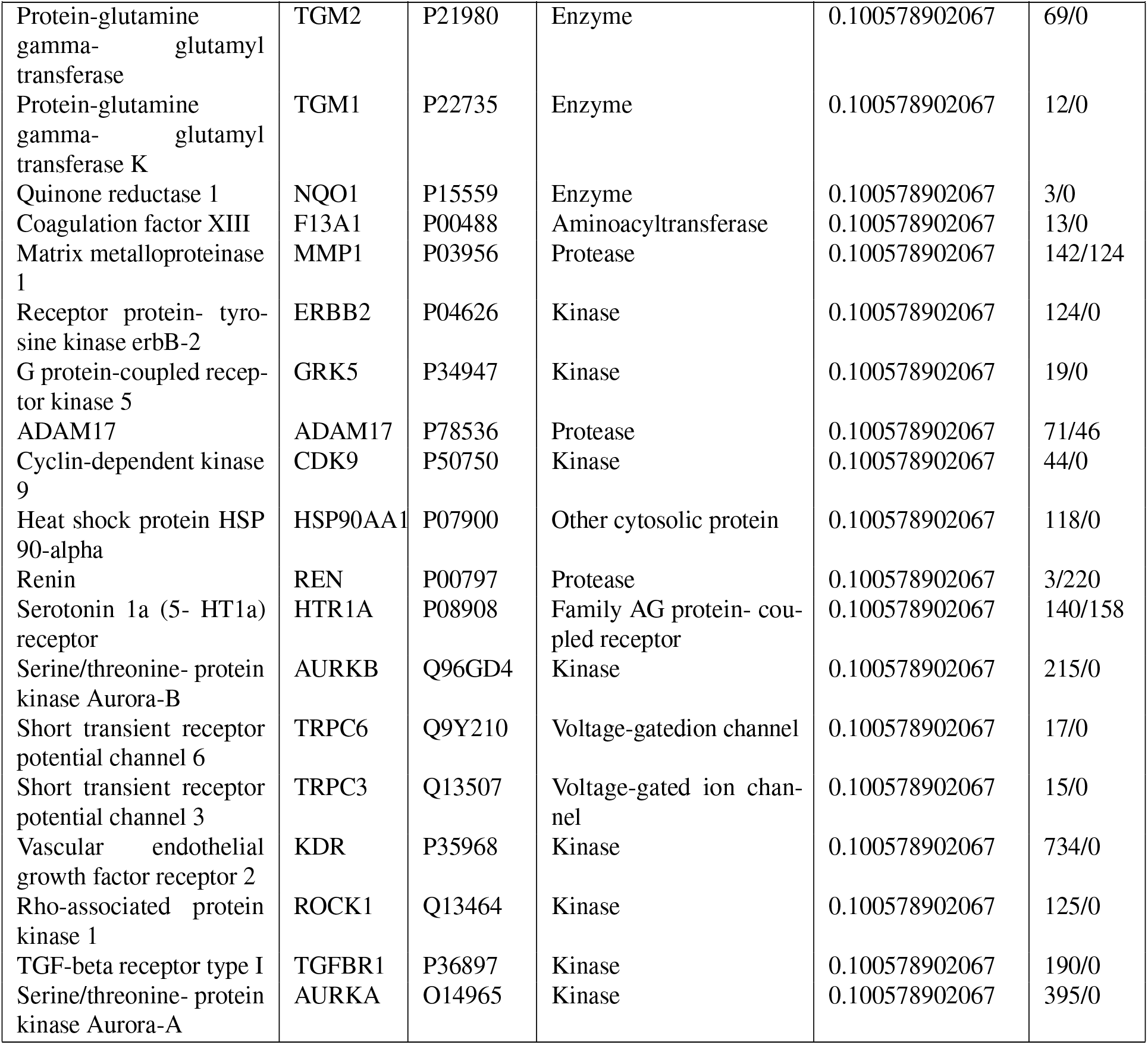
Target Prediction for Rolipram.

**Figure S7.**
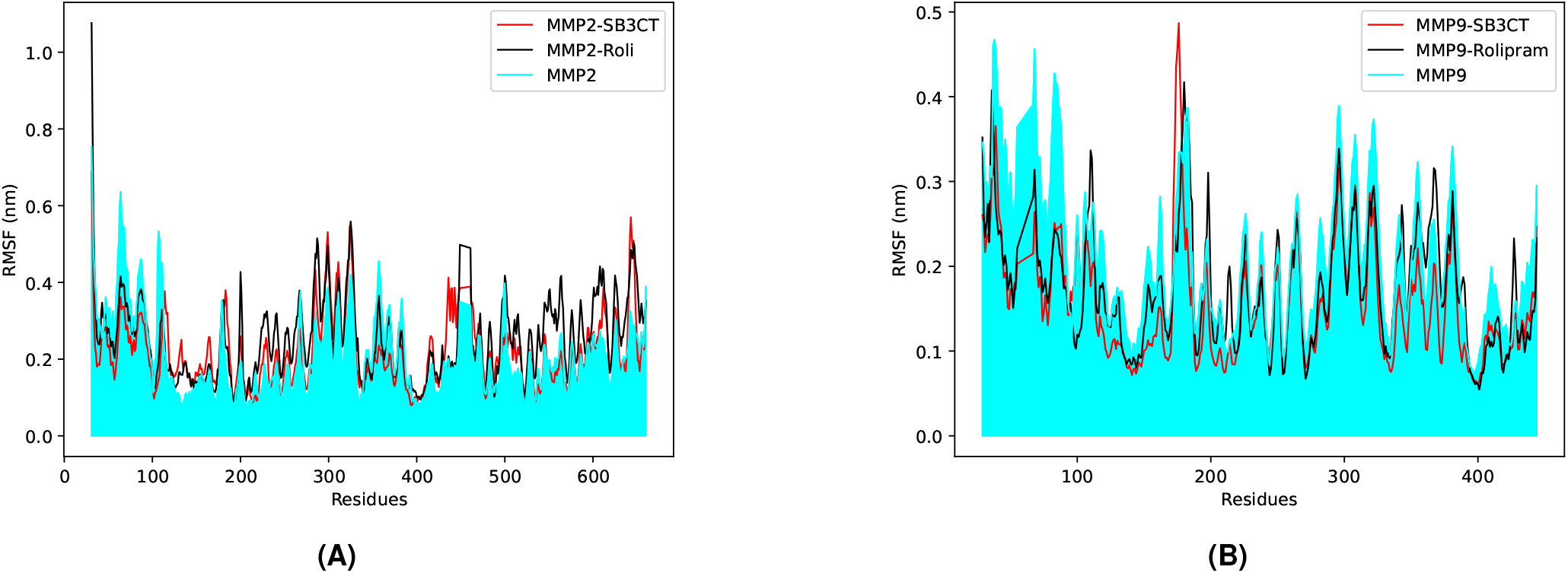
RMSF plots of MMP2-rolipram (A) and MMP2-SB3CT complexes, as well as MMP9-rolipram and MMP9-SB3CT complexes (B), with respect to free MMP2, for the simulation period of 300 *ns*.

**Figure S8.**
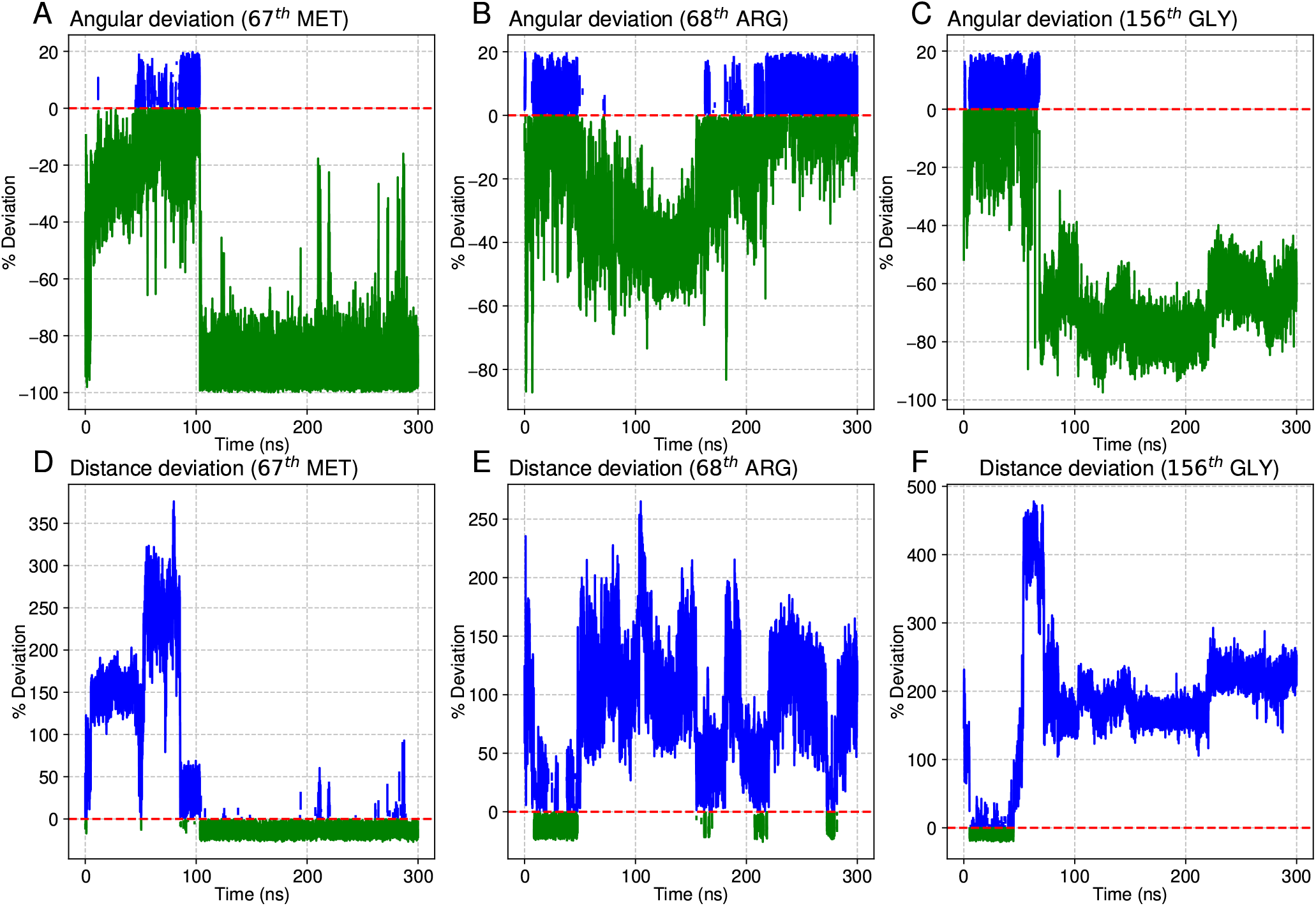
Deviation of angles between hydrogen bond forming atoms from rolipram and from (A) 67^*th*^ MET, (B) 68^*th*^ ARG and (C) 156^*th*^ GLY residues of MMP2. Deviation of the distance between the donor and acceptor atoms of hydrogen bonds formed between rolipram and (D) 67^*th*^ MET, (E) 68^*th*^ ARG and (F) 156^*th*^ GLY residues of MMP2.

**Figure S9.**
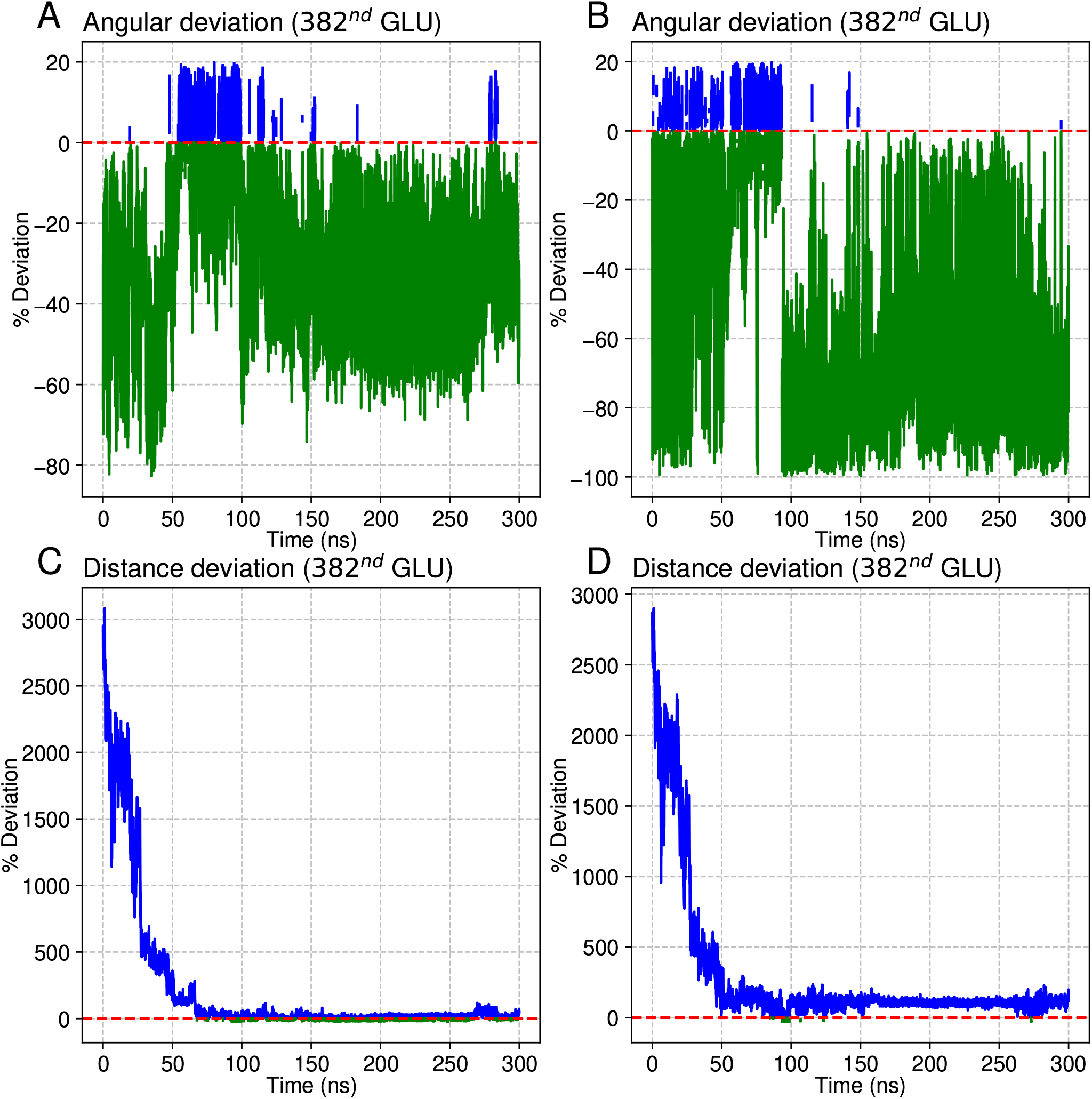
Deviation of angles between hydrogen bond forming atoms from SB3CT and from (A) & (B) 382^*nd*^ GLU residue of MMP2. Deviation of distance between donor and acceptor atoms of hydrogen bonds formed between SB3CT and (C) & (D) 382^*nd*^ GLU residue of MMP2.

**Figure S10.**
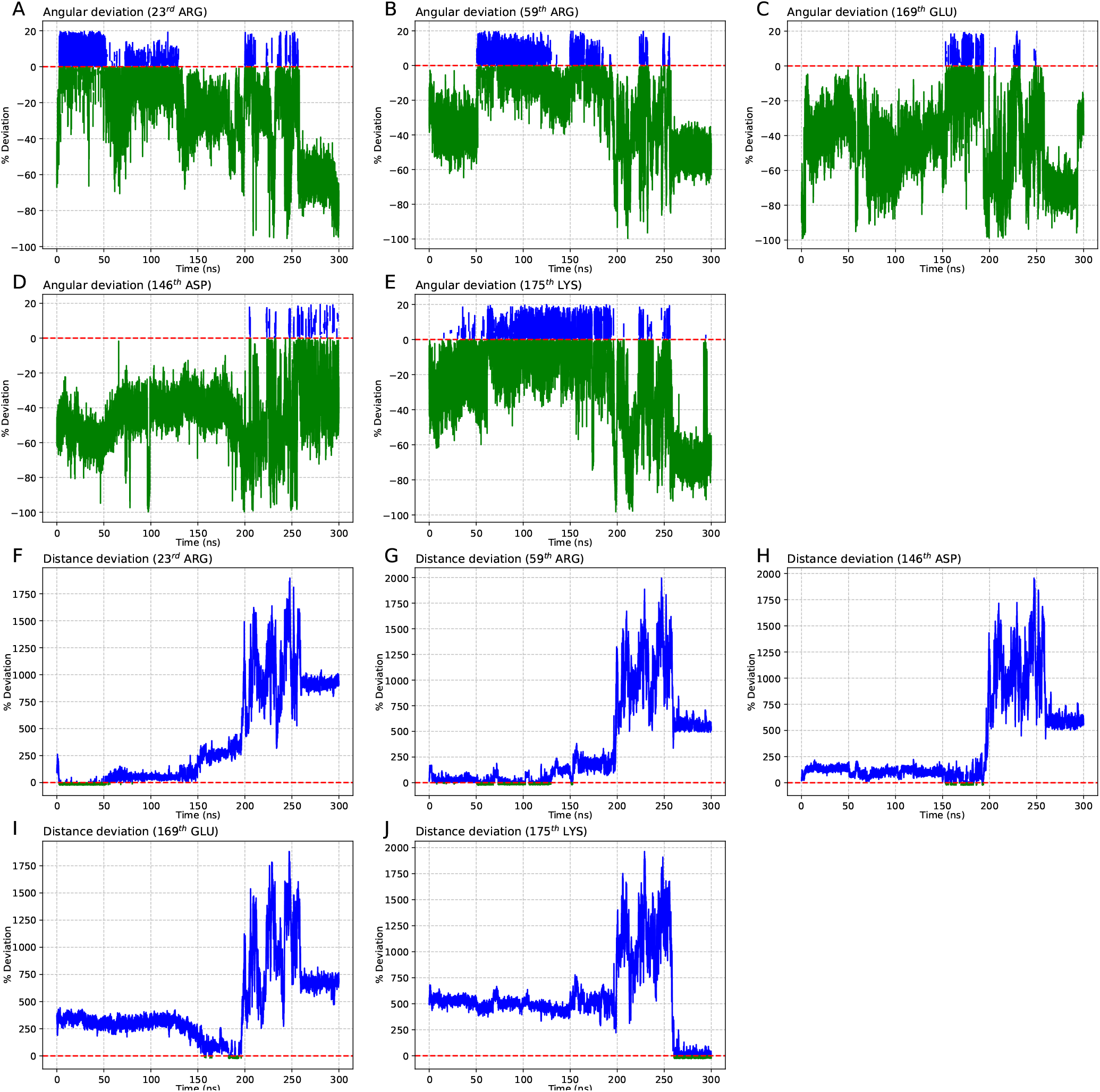
Deviation of angles between hydrogen bond forming atoms from rolipram and from (A) 23^*rd*^ ARG, (B) 59^*th*^ ARG, (C) 169^*th*^ GLU, (D) 146^*th*^ ASP and (E) 175^*th*^ LYS residues of MMP2. Deviation of distance between donor and acceptor atoms of hydrogen bonds formed between rolipram and (F) 23^*rd*^ ARG, (G) 59^*th*^ ARG, (H) 169^*th*^ GLU, (I) 146^*th*^ ASP and (J) 175^*th*^ LYS residues of MMP2.

**Figure S11.**
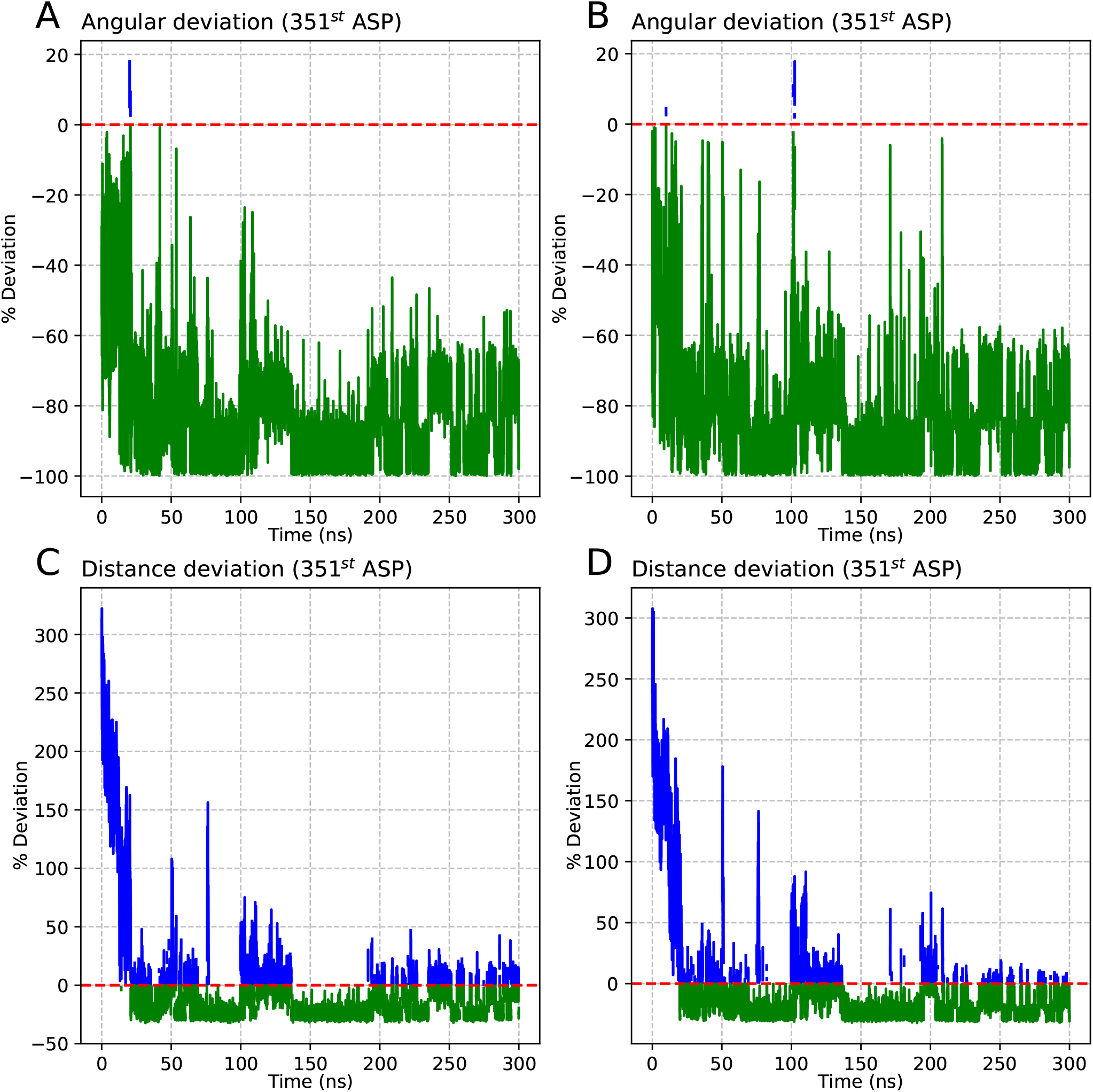
Deviation of angles between hydrogen bond forming atoms from SB3CT and from (A) & (B) 351^*st*^ ASP residue of MMP2. Deviation of distance between donor and acceptor atoms of hydrogen bonds formed between SB3CT and (C) & (D) 351^*st*^ ASP residue of MMP2.

