## Supplementary material for "The Hidden Potential of PDE4 Inhibitor Rolipram: A Multifaceted Examination of its Inhibition of MMP2/9 Reveals Therapeutic Implications": LaTeX Source and figs: MD_paper.bib

@article{mukherjee_pde4_2022,
	title        = {{PDE4} inhibitor eliminates breast cancer stem cells via noncanonical activation of {mTOR}},
	author       = {Mukherjee, Pritha and Bagchi, Arka and Banerjee, Ananya and Roy, Himansu and Bhattacharya, Arijit and Biswas, Arunima and Chatterji, Urmi},
	year         = 2022,
	month        = dec,
	journal      = {Journal of Cellular Biochemistry},
	volume       = 123,
	number       = 12,
	pages        = {1980--1996},
	doi          = {10.1002/jcb.30325},
	issn         = {1097-4644},
	language     = {eng},
	pmid         = 36063486
}
@article{bhaduri_cytotoxic_2022,
	title        = {Cytotoxic activity of nitrogen, sulfur, and oxygen chelated {Pt}({II}) complexes; their {DNA}/{BSA} binding by in vitro and in silico approaches},
	author       = {Bhaduri, Rituparna and Mandal, Saikat and Kumar Tarai, Swarup and Pan, Angana and Mukherjee, Subhajit and Bagchi, Arka and Biswas, Arunima and Ch. Moi, Sankar},
	year         = 2022,
	month        = aug,
	journal      = {Journal of Molecular Liquids},
	volume       = 360,
	pages        = 119529,
	doi          = {10.1016/j.molliq.2022.119529},
	issn         = {0167-7322},
	url          = {https://www.sciencedirect.com/science/article/pii/S0167732222010674},
	urldate      = {2023-06-27},
	language     = {en}
}
@misc{pubchem_rolipram,
	title        = {{N}ational {C}enter for {B}iotechnology {I}nformation. "{P}ub{C}hem {C}ompound {S}ummary for {CID} 5092, {R}olipram" \textit{{P}ub{C}hem},},
	howpublished = {\url{https://pubchem.ncbi.nlm.nih.gov/compound/5092}.},
	note         = {Accessed 20 March, 2024}
}
@article{oboyle_open_2011,
	title        = {Open {Babel}: {An} open chemical toolbox},
	shorttitle   = {Open {Babel}},
	author       = {O'Boyle, Noel M. and Banck, Michael and James, Craig A. and Morley, Chris and Vandermeersch, Tim and Hutchison, Geoffrey R.},
	year         = 2011,
	month        = oct,
	journal      = {Journal of Cheminformatics},
	volume       = 3,
	number       = 1,
	pages        = 33,
	doi          = {10.1186/1758-2946-3-33},
	issn         = {1758-2946},
	url          = {https://doi.org/10.1186/1758-2946-3-33},
	urldate      = {2023-09-09}
}
@article{lindahl_molecular_2008,
	title        = {Molecular dynamics simulations},
	author       = {Lindahl, Erik R.},
	year         = 2008,
	journal      = {Methods in Molecular Biology (Clifton, N.J.)},
	volume       = 443,
	pages        = {3--23},
	doi          = {10.1007/978-1-59745-177-2\_1},
	issn         = {1064-3745},
	language     = {eng},
	pmid         = 18446279
}
@misc{arka:github,
	title        = {{G}ROMACS {F}iles},
	author       = {Bagchi, A.},
	year         = 2023,
	journal      = {GitHub repository},
	publisher    = {GitHub},
	howpublished = {\url{https://github.com/arkabagchi1993/GROMACS\_Files}}
}
@article{zoete_swissparam_2011,
	title        = {{SwissParam}: a fast force field generation tool for small organic molecules},
	shorttitle   = {{SwissParam}},
	author       = {Zoete, Vincent and Cuendet, Michel A. and Grosdidier, Aur\'{e}lien and Michielin, Olivier},
	year         = 2011,
	month        = aug,
	journal      = {Journal of Computational Chemistry},
	volume       = 32,
	number       = 11,
	pages        = {2359--2368},
	doi          = {10.1002/jcc.21816},
	issn         = {1096-987X},
	language     = {eng},
	pmid         = 21541964
}
@misc{tresanco_amdock_2023,
	title        = {{AMDock}: {Assisted} {Molecular} {Docking} with {Autodock4} and {Autodock} {Vina}},
	shorttitle   = {{AMDock}},
	author       = {Tresanco, Mario Sergio Vald\'{e}s},
	year         = 2023,
	month        = sep,
	url          = {https://github.com/Valdes-Tresanco-MS/AMDock},
	urldate      = {2023-09-29},
	copyright    = {GPL-3.0},
	note         = {original-date: 2018-02-20T16:41:39Z}
}
@misc{pubchem_sb3ct,
	title        = {{N}ational {C}enter for {B}iotechnology {I}nformation. "{P}ub{C}hem {C}ompound {S}ummary for {CID} 9883002, 2-[(4-{P}henoxyphenyl)sulfonylmethyl]thiirane" \textit{{P}ub{C}hem},},
	howpublished = {\url{https://pubchem.ncbi.nlm.nih.gov/compound/2-\_4-Phenoxyphenyl\_sulfonylmethyl\_thiirane}.},
	year         = 2024
}
@article{rudra_melatonin_2013,
	title        = {Melatonin inhibits matrix metalloproteinase-9 activity by binding to its active site},
	author       = {Rudra, Deep Sankar and Pal, Uttam and Maiti, Nakul Chandra and Reiter, Russel J. and Swarnakar, Snehasikta},
	year         = 2013,
	journal      = {Journal of Pineal Research},
	volume       = 54,
	number       = 4,
	pages        = {398--405},
	doi          = {10.1111/jpi.12034},
	issn         = {1600-079X},
	url          = {https://onlinelibrary.wiley.com/doi/abs/10.1111/jpi.12034},
	urldate      = {2024-01-05},
	copyright    = {\textcopyright{} 2012 John Wiley \& Sons A/S. Published by Blackwell Publishing Ltd},
	note         = {\_eprint: https://onlinelibrary.wiley.com/doi/pdf/10.1111/jpi.12034},
	language     = {en}
}
@article{toth_assessment_2012,
	title        = {Assessment of gelatinases ({MMP}-2 and {MMP}-9) by gelatin zymography},
	author       = {Toth, Marta and Sohail, Anjum and Fridman, Rafael},
	year         = 2012,
	journal      = {Methods in Molecular Biology (Clifton, N.J.)},
	volume       = 878,
	pages        = {121--135},
	doi          = {10.1007/978-1-61779-854-2\_8},
	issn         = {1940-6029},
	language     = {eng},
	pmid         = 22674130
}
@article{abraham_gromacs_2015,
	title        = {{GROMACS}: {High} performance molecular simulations through multi-level parallelism from laptops to supercomputers},
	shorttitle   = {{GROMACS}},
	author       = {Abraham, Mark James and Murtola, Teemu and Schulz, Roland and P\'{a}ll, Szil\'{a}rd and Smith, Jeremy C. and Hess, Berk and Lindahl, Erik},
	year         = 2015,
	month        = sep,
	journal      = {SoftwareX},
	volume       = {1-2},
	pages        = {19--25},
	doi          = {10.1016/j.softx.2015.06.001},
	issn         = {2352-7110},
	url          = {https://www.sciencedirect.com/science/article/pii/S2352711015000059},
	urldate      = {2024-01-05}
}
@inproceedings{pall_tackling_2015,
	title        = {Tackling {Exascale} {Software} {Challenges} in {Molecular} {Dynamics} {Simulations} with {GROMACS}},
	author       = {P\'{a}ll, Szil\'{a}rd and Abraham, Mark James and Kutzner, Carsten and Hess, Berk and Lindahl, Erik},
	year         = 2015,
	booktitle    = {Solving {Software} {Challenges} for {Exascale}},
	publisher    = {Springer International Publishing},
	address      = {Cham},
	series       = {Lecture {Notes} in {Computer} {Science}},
	pages        = {3--27},
	doi          = {10.1007/978-3-319-15976-8\_1},
	isbn         = {978-3-319-15976-8},
	language     = {en},
	editor       = {Markidis, Stefano and Laure, Erwin}
}
@article{chen_sonic_2013,
	title        = {Sonic hedgehog signaling pathway induces cell migration and invasion through focal adhesion kinase/{AKT} signaling-mediated activation of matrix metalloproteinase ({MMP})-2 and {MMP}-9 in liver cancer},
	author       = {Chen, Jing-Song and Huang, Xiao-hui and Wang, Qian and Huang, Jiong-Qiang and Zhang, Long-juan and Chen, Xi-Lin and Lei, Jian and Cheng, Zhi-Xiang},
	year         = 2013,
	month        = jan,
	journal      = {Carcinogenesis},
	volume       = 34,
	number       = 1,
	pages        = {10--19},
	doi          = {10.1093/carcin/bgs274},
	issn         = {0143-3334},
	url          = {https://doi.org/10.1093/carcin/bgs274},
	urldate      = {2024-01-05}
}
@article{vihinen_matrix_2005,
	title        = {Matrix metalloproteinases as therapeutic targets in cancer},
	author       = {Vihinen, Pia and Ala-aho, Risto and K\"{a}h\"{a}ri, Veli-Matti},
	year         = 2005,
	month        = may,
	journal      = {Current Cancer Drug Targets},
	volume       = 5,
	number       = 3,
	pages        = {203--220},
	doi          = {10.2174/1568009053765799},
	issn         = {1568-0096},
	language     = {eng},
	pmid         = 15892620
}
@article{lemkul_proteins_2019,
	title        = {From {Proteins} to {Perturbed} {Hamiltonians}: {A} {Suite} of {Tutorials} for the {GROMACS}-2018 {Molecular} {Simulation} {Package} [{Article} v1.0]},
	shorttitle   = {From {Proteins} to {Perturbed} {Hamiltonians}},
	author       = {Lemkul, Justin A.},
	year         = 2019,
	journal      = {Living Journal of Computational Molecular Science},
	volume       = 1,
	number       = 1,
	pages        = {5068--5068},
	doi          = {10.33011/livecoms.1.1.5068},
	issn         = {2575-6524},
	url          = {https://livecomsjournal.org/index.php/livecoms/article/view/v1i1e5068},
	urldate      = {2024-01-05},
	copyright    = {Copyright (c) 2018},
	note         = {Number: 1},
	language     = {en}
}
@article{mackerell_all-atom_1998,
	title        = {All-atom empirical potential for molecular modeling and dynamics studies of proteins},
	author       = {MacKerell, A. D. and Bashford, D. and Bellott, M. and Dunbrack, R. L. and Evanseck, J. D. and Field, M. J. and Fischer, S. and Gao, J. and Guo, H. and Ha, S. and Joseph-McCarthy, D. and Kuchnir, L. and Kuczera, K. and Lau, F. T. and Mattos, C. and Michnick, S. and Ngo, T. and Nguyen, D. T. and Prodhom, B. and Reiher, W. E. and Roux, B. and Schlenkrich, M. and Smith, J. C. and Stote, R. and Straub, J. and Watanabe, M. and Wi\'{o}rkiewicz-Kuczera, J. and Yin, D. and Karplus, M.},
	year         = 1998,
	month        = apr,
	journal      = {The Journal of Physical Chemistry. B},
	volume       = 102,
	number       = 18,
	pages        = {3586--3616},
	doi          = {10.1021/jp973084f},
	issn         = {1520-6106},
	language     = {eng},
	pmid         = 24889800
}
@article{michaud-agrawal_mdanalysis_2011,
	title        = {{MDAnalysis}: a toolkit for the analysis of molecular dynamics simulations},
	shorttitle   = {{MDAnalysis}},
	author       = {Michaud-Agrawal, Naveen and Denning, Elizabeth J. and Woolf, Thomas B. and Beckstein, Oliver},
	year         = 2011,
	month        = jul,
	journal      = {Journal of Computational Chemistry},
	volume       = 32,
	number       = 10,
	pages        = {2319--2327},
	doi          = {10.1002/jcc.21787},
	issn         = {1096-987X},
	language     = {eng},
	pmid         = 21500218,
	pmcid        = {PMC3144279}
}
@article{luzar_hydrogen-bond_1996,
	title        = {Hydrogen-bond kinetics in liquid water},
	author       = {Luzar, Alenka and Chandler, David},
	year         = 1996,
	month        = jan,
	journal      = {Nature},
	volume       = 379,
	number       = 6560,
	pages        = {55--57},
	doi          = {10.1038/379055a0},
	issn         = {1476-4687},
	url          = {https://www.nature.com/articles/379055a0},
	urldate      = {2024-01-05},
	copyright    = {1996 Springer Nature Limited},
	note         = {Number: 6560 Publisher: Nature Publishing Group},
	language     = {en}
}
@article{luzar_resolving_2000,
	title        = {Resolving the hydrogen bond dynamics conundrum},
	author       = {Luzar, Alenka},
	year         = 2000,
	month        = dec,
	journal      = {The Journal of Chemical Physics},
	volume       = 113,
	number       = 23,
	pages        = {10663--10675},
	doi          = {10.1063/1.1320826},
	issn         = {0021-9606},
	url          = {https://doi.org/10.1063/1.1320826},
	urldate      = {2024-01-05}
}
@article{almutairi_matrix_2023,
	title        = {Matrix {Metalloproteinases} {Inhibitors} in {Cancer} {Treatment}: {An} {Updated} {Review} (2013–2023)},
	shorttitle   = {Matrix {Metalloproteinases} {Inhibitors} in {Cancer} {Treatment}},
	author       = {Almutairi, Shriefa and Kalloush, Hanin Moh'd and Manoon, Nour A. and Bardaweel, Sanaa K.},
	year         = 2023,
	month        = jul,
	journal      = {Molecules},
	volume       = 28,
	number       = 14,
	pages        = 5567,
	doi          = {10.3390/molecules28145567},
	issn         = {1420-3049},
	url          = {https://www.ncbi.nlm.nih.gov/pmc/articles/PMC10384300/},
	urldate      = {2024-03-05},
	pmid         = 37513440,
	pmcid        = {PMC10384300}
}
@article{freije_matrix_2003,
	title        = {Matrix metalloproteinases and tumor progression},
	author       = {Freije, Jos\'{e} M. P. and Balb\'{\i}n, Milagros and Pend\'{a}s, Alberto M. and S\'{a}nchez, Luis M. and Puente, Xose S. and L\'{o}pez-Ot\'{\i}n, Carlos},
	year         = 2003,
	journal      = {Advances in Experimental Medicine and Biology},
	volume       = 532,
	pages        = {91--107},
	doi          = {10.1007/978-1-4615-0081-0\_9},
	issn         = {0065-2598},
	language     = {eng},
	pmid         = 12908552
}
@article{ahmad_role_2018,
	title        = {Role of {MMP}-2, {MMP}-9 and {VEGF} as serum biomarker in early prognosis of renal cell carcinoma},
	author       = {Ahmad, S. and Singh, V. and Sinha, R. J. and Srivastava, A. and Mandhani, A.},
	year         = 2018,
	month        = dec,
	journal      = {African Journal of Urology},
	volume       = 24,
	number       = 4,
	pages        = {255--263},
	doi          = {10.1016/j.afju.2018.09.003},
	issn         = {1110-5704},
	url          = {https://www.sciencedirect.com/science/article/pii/S1110570418301231},
	urldate      = {2024-03-05}
}
@article{benitha_evaluation_nodate,
	title        = {Evaluation of {Serum} {Levels} of {Matrix} {MetalloProteinase}-9 ({MMP}-9) in {Oral} {Squamous} {Cell} {Carcinoma} and {Its} {Clinicopathological} {Correlation}},
	author       = {Benitha, Georgia and Ramani, Pratibha and Jayaraman, Selvaraj and R, Abilasha and Ramalingam, Karthikeyan and Krishnan, Murugesan},
	year         = 2023,
	journal      = {Cureus},
	volume       = 15,
	number       = 2,
	pages        = {e34954},
	doi          = {10.7759/cureus.34954},
	issn         = {2168-8184},
	url          = {https://www.ncbi.nlm.nih.gov/pmc/articles/PMC10018444/},
	urldate      = {2024-03-05},
	pmid         = 36938194,
	pmcid        = {PMC10018444}
}
@article{Bagchi2024.03.03.583221,
	title        = {PDE4 inhibitor Rolipram represses hedgehog signaling via ubiquitin-mediated proteolysis of GLI transcription factors to regress breast cancer},
	author       = {Arka Bagchi and Anuran Bhattacharya and Urmi Chatterji and ARUNIMA BISWAS},
	year         = 2024,
	journal      = {bioRxiv},
	publisher    = {Cold Spring Harbor Laboratory},
	doi          = {10.1101/2024.03.03.583221},
	url          = {https://www.biorxiv.org/content/early/2024/03/06/2024.03.03.583221},
	elocation-id = {2024.03.03.583221},
	eprint       = {https://www.biorxiv.org/content/early/2024/03/06/2024.03.03.583221.full.pdf}
}

@article{li_relationship_2017,
	title = {The relationship between {MMP}-2 and {MMP}-9 expression levels with breast cancer incidence and prognosis},
	volume = {14},
	issn = {1792-1074},
	doi = {10.3892/ol.2017.6924},
	language = {eng},
	number = {5},
	journal = {Oncology Letters},
	author = {Li, Hai and Qiu, Zhenwei and Li, Feng and Wang, Chunlei},
	month = nov,
	year = {2017},
	pmid = {29113219},
	pmcid = {PMC5661385},
	keywords = {breast cancer, matrix metalloproteinase-2, matrix metalloproteinase-9, prognosis},
	pages = {5865--5870},
}


@article{jiang_prognostic_2021,
	title = {Prognostic values of tumoral {MMP2} and {MMP9} overexpression in breast cancer: a systematic review and meta-analysis},
	volume = {21},
	issn = {1471-2407},
	url = {https://doi.org/10.1186/s12885-021-07860-2},
	doi = {10.1186/s12885-021-07860-2},
	number = {1},
	journal = {BMC Cancer},
	author = {Jiang, Hanfang and Li, Huiping},
	month = feb,
	year = {2021},
	pages = {149},
}


@article{muniz-bongers_mmp2_nodate,
	title = {{MMP2} and {TLRs} modulate immune responses in the tumor microenvironment},
	volume = {6},
    year = {2021},
	issn = {2379-3708},
	url = {https://www.ncbi.nlm.nih.gov/pmc/articles/PMC8262464/},
	doi = {10.1172/jci.insight.144913},
	number = {12},
	urldate = {2024-03-27},
	journal = {JCI Insight},
	author = {Muniz-Bongers, Luciana R. and McClain, Christopher B. and Saxena, Mansi and Bongers, Gerold and Merad, Miriam and Bhardwaj, Nina},
	pmid = {34032639},
	pmcid = {PMC8262464},
	pages = {e144913},
}


@article{leifler_inflammation_2013,
	title = {Inflammation {Induced} by {MMP}-9 {Enhances} {Tumor} {Regression} of {Experimental} {Breast} {Cancer}},
	volume = {190},
	issn = {0022-1767},
	url = {https://www.ncbi.nlm.nih.gov/pmc/articles/PMC3619527/},
	doi = {10.4049/jimmunol.1202610},
	number = {8},
	urldate = {2024-03-27},
	journal = {The Journal of Immunology Author Choice},
	author = {Leifler, Karin Söderlund and Svensson, Susanne and Abrahamsson, Annelie and Bendrik, Christina and Robertson, Jennifer and Gauldie, Jack and Olsson, Anna-Karin and Dabrosin, Charlotta},
	month = apr,
	year = {2013},
	pmid = {23509357},
	pmcid = {PMC3619527},
	pages = {4420--4430},
	file = {PubMed Central Full Text PDF:C\:\\Users\\Hp\\Zotero\\storage\\K4ZUV6KS\\Leifler et al. - 2013 - Inflammation Induced by MMP-9 Enhances Tumor Regre.pdf:application/pdf},
}
