## Supplementary figures and images for "The Hidden Potential of PDE4 Inhibitor Rolipram: A Multifaceted Examination of its Inhibition of MMP2/9 Reveals Therapeutic Implications"

### Fig1.jpg

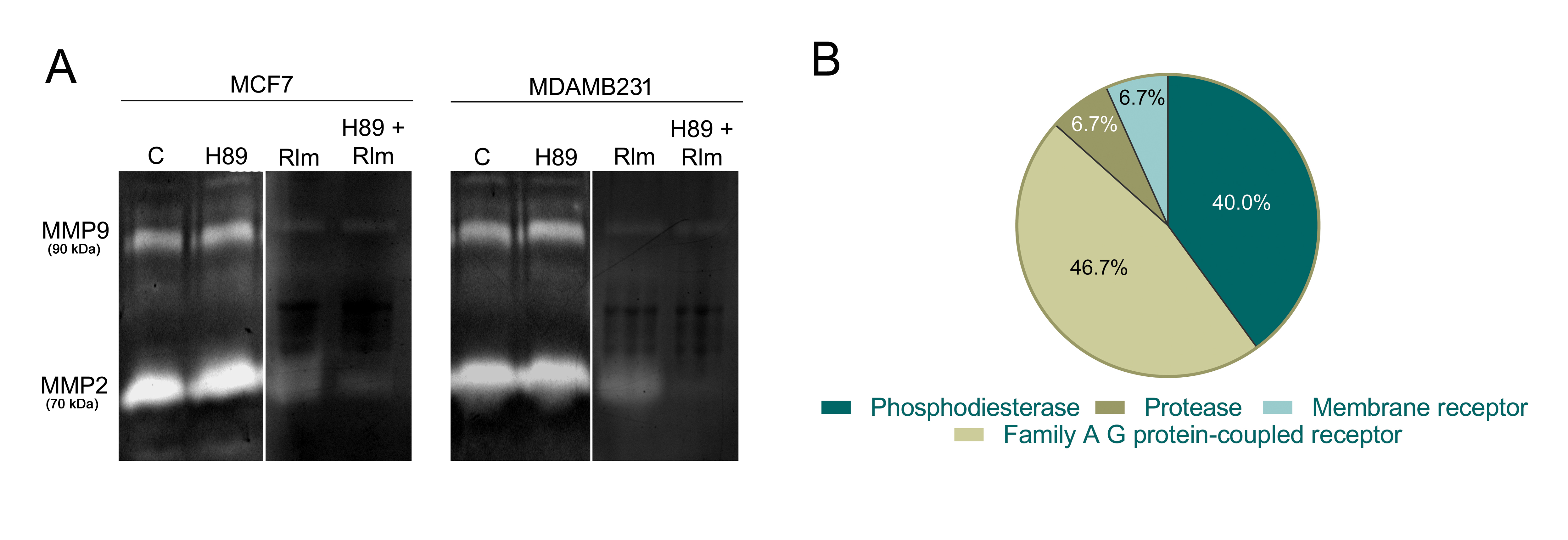

### Fig2.jpg

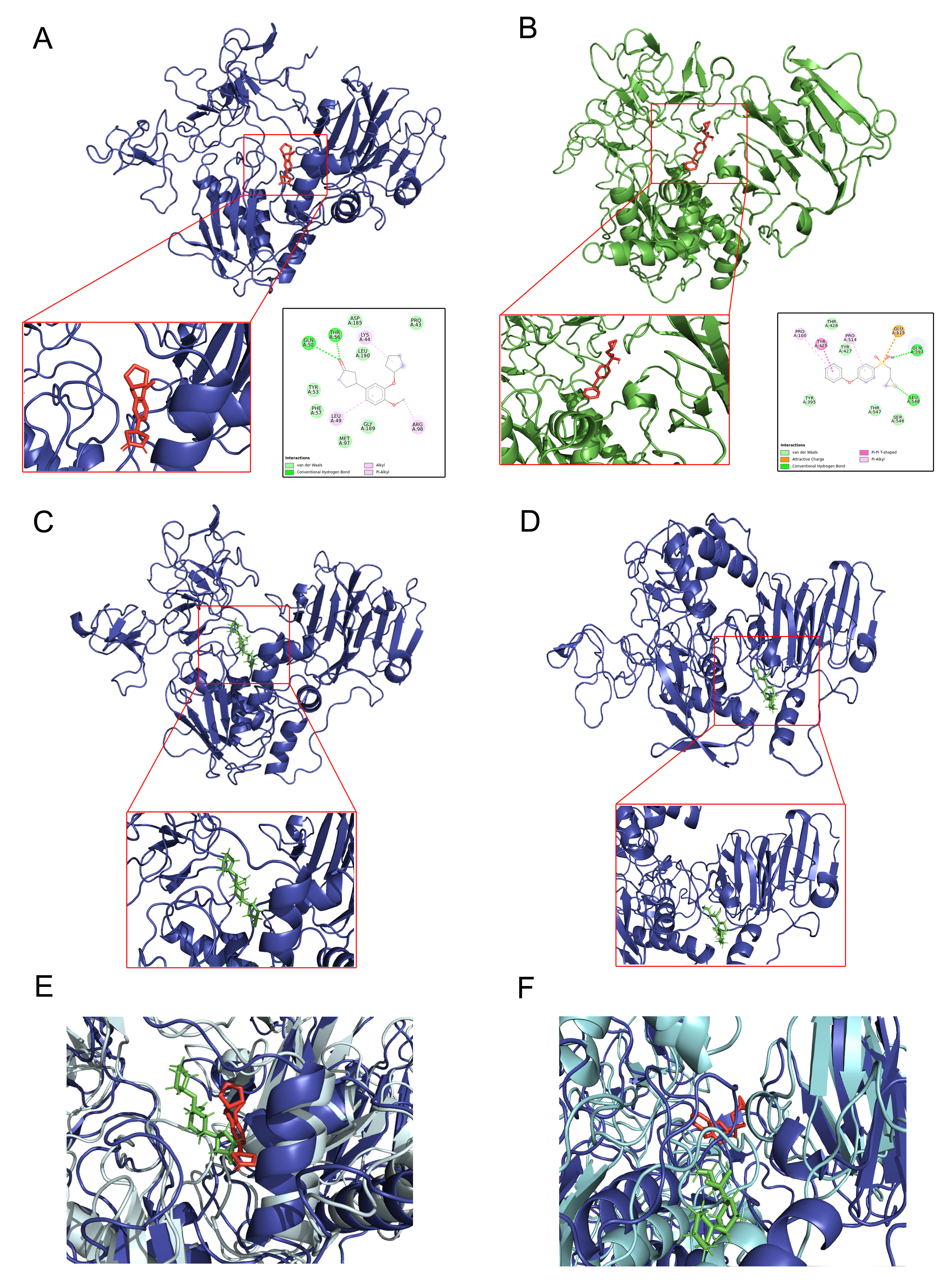

### Fig4.jpg

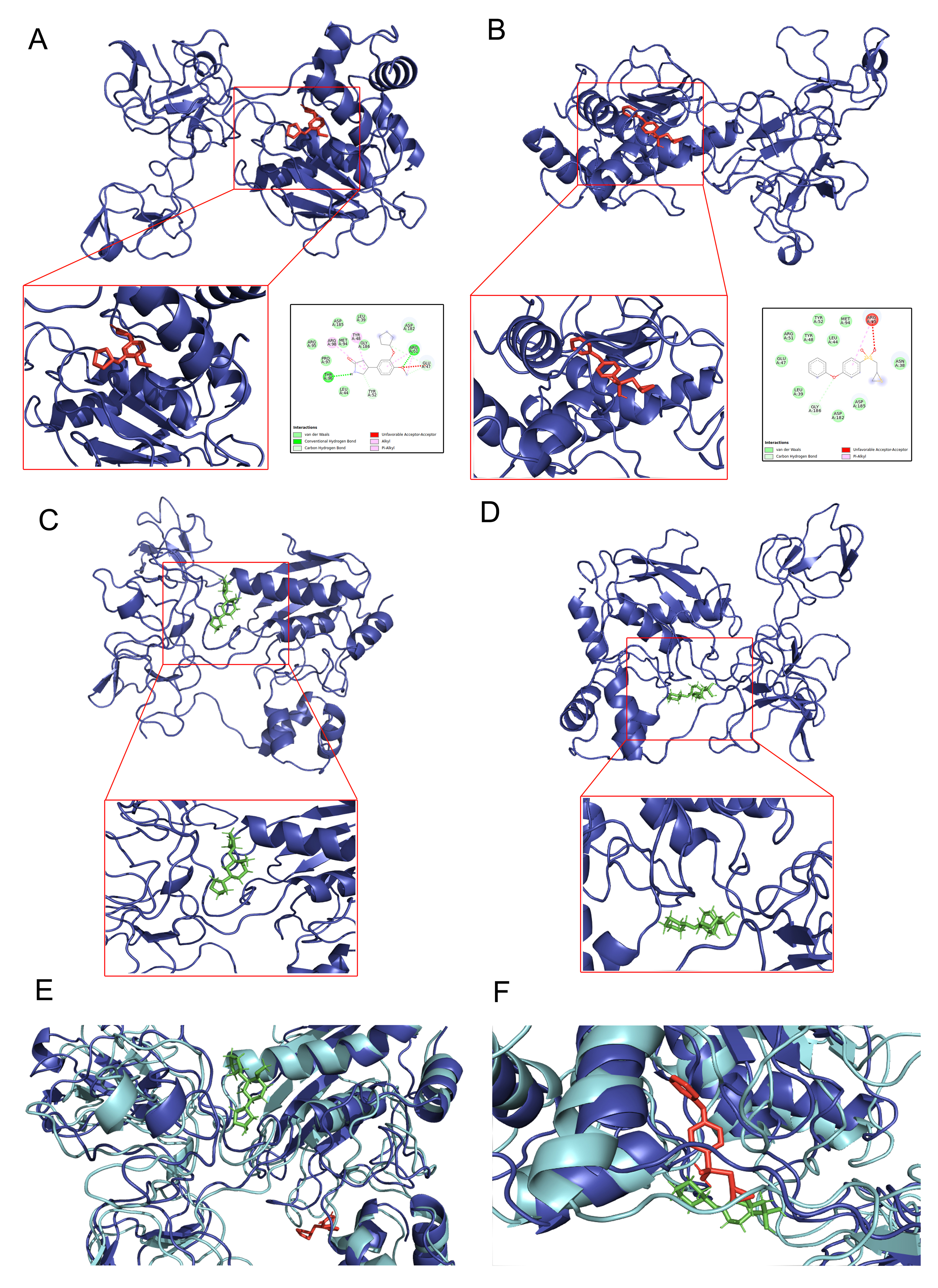

### Fig6.jpg

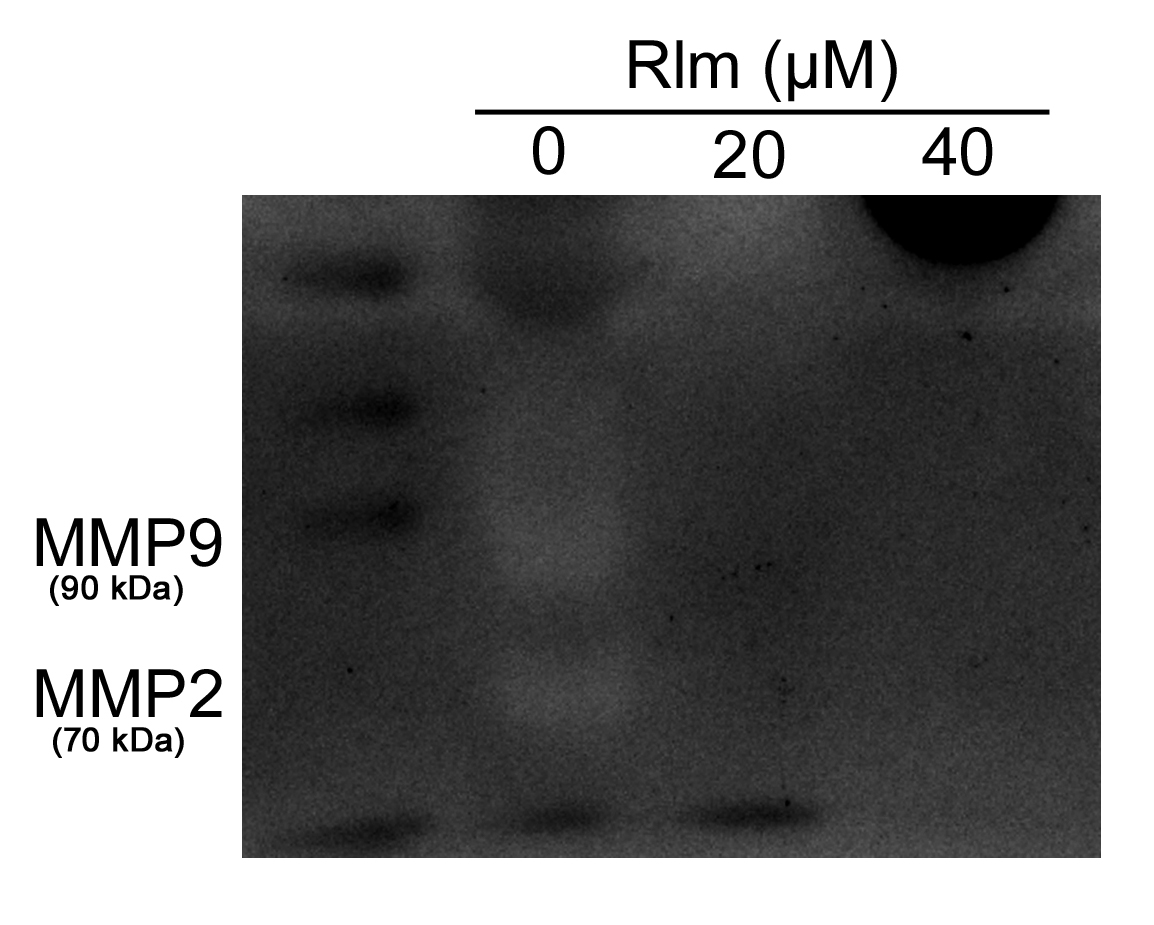
